# An interkinetic envelope surrounds chromosomes between meiosis I and II in C. elegans oocytes

**DOI:** 10.1101/2024.10.19.619195

**Authors:** Layla El Mossadeq, Laura Bellutti, Rémi Le Borgne, Julie C. Canman, Lionel Pintard, Jean-Marc Verbavatz, Peter Askjaer, Julien Dumont

## Abstract

At the end of cell division, the nuclear envelope reassembles around the decondensing chromosomes. Female meiosis culminates in two consecutive cell divisions of the oocyte, meiosis I and II, which are separated by a brief transition phase known as interkinesis. Due to the absence of chromosome decondensation and the suppression of genome replication during interkinesis, it has been widely assumed that the nuclear envelope does not reassemble between meiosis I and II. By analyzing interkinesis in *C. elegans* oocytes, we instead show that an atypical structure made of two lipid bilayers, which we termed the interkinetic envelope, surrounds the surface of the segregating chromosomes. The interkinetic envelope shares common features with the nuclear envelope but also exhibits specific characteristics that distinguish it, including its lack of continuity with the endoplasmic reticulum, unique protein composition, assembly mechanism, and function in chromosome segregation. These distinct attributes collectively define the interkinetic envelope as a unique and specialized structure that has been previously overlooked.

## INTRODUCTION

The nuclear envelope delineates the nucleus in all eukaryotic cells. The nuclear envelope is comprised of two lipid bilayers, which form the inner nuclear membrane (INM) in contact with chromatin, and the outer nuclear membrane (ONM) facing the cytoplasm (Hetzer, 2010). Distinct protein compositions characterize the two layers of the nuclear envelope. The INM, lined by the nuclear lamina, faces the nucleoplasmic compartment, and features a unique set of proteins, including the LAP2, Emerin, and MAN1 (LEM)-domain integral membrane proteins (Ungricht and Kutay, 2015). The ONM continuously connects to the endoplasmic reticulum (ER) and shares both composition and function with the ER (Deolal et al., 2024). While the primary role of the nuclear envelope is to separate the genome and nucleoplasmic space from the cytoplasm, there are specific points of contact and communication between these two compartments. First, the INM is fused with the ONM at designated sites where multisubunit macromolecular complexes, known as nuclear pore complexes (NPCs), assemble and facilitate nucleocytoplasmic transport across the nuclear envelope (De Magistris and Antonin, 2018; Ungricht and Kutay, 2017). Second, the linker of nucleoskeleton and cytoskeleton (LINC) complex, a highly conserved 6:6 heterohexameric bridge spanning the nuclear envelope, serves to physically connect chromatin and the nuclear lamina to the cytoskeleton (McGillivary et al., 2023).

In organisms undergoing semi-open or open mitosis, the transition from interphase to mitosis (M-phase) is marked by nuclear envelope breakdown (NEBD) and chromosome condensation (Boettcher and Barral, 2013). Following cell division and the segregation of sister chromatids, the nuclear envelope must reassemble around the decondensing chromatids to separate the genome from the cytoplasmic environment. Thus, cycles of NEBD and chromosome condensation, followed by nuclear envelope reassembly around decondensing chromatids, accompany successive cell divisions in most tissues and cell types. A notable deviation from this stereotypical sequence of events occurs during oogenesis. This process, responsible for producing haploid female gametes, culminates in two consecutive cell divisions of the oocyte, known as meiosis I and II (Dumont and Desai, 2012; Mullen et al., 2019; Ohkura, 2015; Severson et al., 2016). During meiosis I, recombined homologous chromosome pairs are segregated into two chromosome sets. One set is directed for elimination into the first polar body (hereafter referred to as the PB chromosomal set), while the second set almost immediately proceeds to meiosis II (hereafter referred to as the MII chromosomal set), following a very brief transition phase termed interkinesis. A remarkable feature of interkinesis is the apparent lack of chromosome decondensation preceding entry into meiosis II and the segregation of sister chromatids (Nakajo et al., 2000). The absence of chromosome decondensation at this stage is coupled with suppression of genome replication, which normally occurs after exit from M-phase, and which is essential in this specific context for generating haploid oocytes (Furuno et al., 1994). Hence, although interkinesis occurs between two M-phases, it is not classified as a typical interphase. In this context, the status of the nuclear envelope during interkinesis remains notably ambiguous (Gerhart et al., 1984; Lenart and Ellenberg, 2003; Nakajo et al., 2000; Nebreda and Ferby, 2000). As interkinesis occurs between two M-phases, one would anticipate nuclear envelope reassembly in the oocyte at this stage. Yet, the apparent suppression of most interphase events and the scarcity of reports on the presence of a canonical nuclear envelope surrounding oocyte chromosomes during interkinesis in any species has led to controversy over its actual existence (Penfield et al., 2020).

By combining light and electron microscopy, we probed nuclear envelope reassembly in oocytes of the nematode *Caenorhabditis elegans* during interkinesis. We found that a double membrane, superficially reminiscent of the nuclear envelope, progressively assembles at the surface of the segregating chromosomes during anaphase/telophase I. This structure is transient and disassembles rapidly upon entry into meiosis II. Furthermore, examination of the ultrastructure, protein composition, and function of this double membrane in *C. elegans* oocytes revealed distinctive structural, compositional, and functional features that set it apart from a typical nuclear envelope. We thus named this novel organelle the interkinetic envelope.

## RESULTS

### An interkinetic envelope forms on the surface of both chromosomal sets between meiosis I and II in oocytes

To determine if a nuclear envelope reassembles during the short interkinetic transition phase between meiosis I and II in *C. elegans* oocytes, we analyzed the 3- dimensional organization of membranes around chromosomes during anaphase I/interkinesis by correlative light and serial block-face scanning electron microscopy (SBF-SEM) (Lachat et al., 2022). Fertilized oocytes expressing green fluorescent protein (GFP)-tagged tubulin and mCherry-tagged H2B were imaged *ex utero* using a spinning disk microscope until they reached mid-anaphase I or mid/late-interkinesis (Fig. 1 A). They were then fixed chemically and processed for SBF-SEM. 30 nm-thick sections were automatically cut and imaged throughout the two sets of segregating chromosomes, and a slab of each stage oocyte including both sets of chromosomes was reconstructed (Fig. 1 B and Video 1).

**Figure 1:**
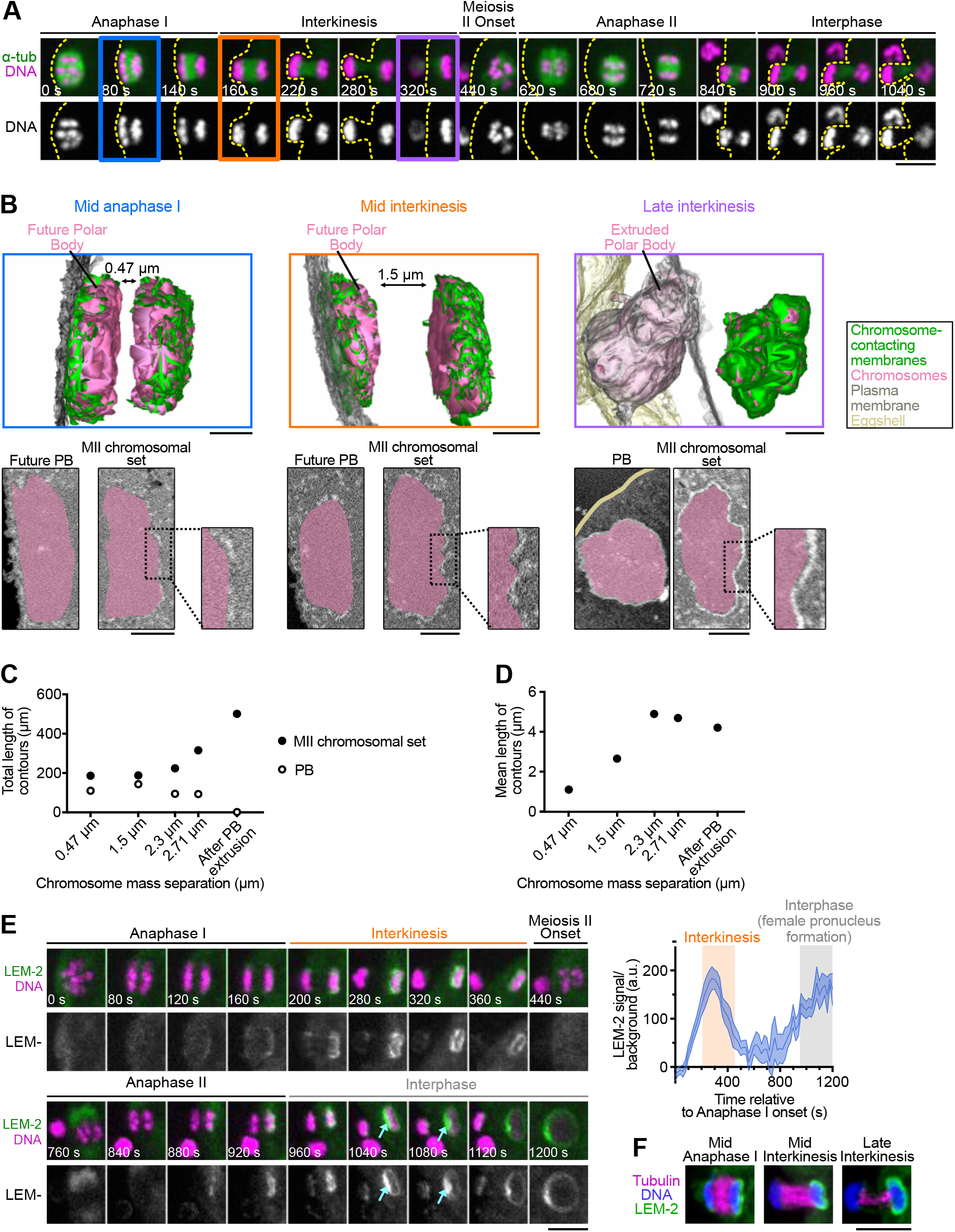
The interkinetic envelope forms between meiosis I and II in the *C. elegans* oocyte. **(A)** Representative time-lapse images of GFP::TBA-2^α-tubulin^ (green) and mCherry::HIS-11^H2B^ (magenta)-expressing oocytes during meiosis I and II (n= 7). Timings indicated at the bottom left corners of images are from anaphase I onset. The specific meiotic stages used for electron microscopy are highlighted in blue (mid-anaphase), orange (mid interkinesis) and purple (late interkinesis). Scale bar, 5 μm. **(B)** 3-dimensional reconstructions centered on chromosomes of a mid-anaphase I (left, n= 1), a mid-interkinesis (center, n= 1) and a late-interkinesis (right, n= 1) oocytes acquired by SBF-SEM. Chromosomes in magenta, membranes in contact with chromosomes in green, plasma membrane in gray, and eggshell in gold. Each reconstruction is accompanied (bottom) by a 2-dimensional single section showing each chromosome set in magenta, and a magnification of a region of interest (ROI) of the MII chromosomal set. Scale bar, 1 µm. **(C, D)** Quantifications of the total (C) and mean (D) length of membrane contours from five reconstructed oocytes represented according to the distance between the segregating chromosomal sets (chromosome mass separation) for the polar body chromosomal set (empty dots) and for the MII chromosomal set (solid dots). **(E)** Left: Representative time-lapse images of GFP::LEM-2^LEMD2/3^ and mCherry::H2B expressing oocytes (n= 9) during meiosis I (top) and meiosis II (bottom). Timing relative to anaphase I onset is indicated at the bottom left corner of each image. The cyan arrows indicate the GFP::LEM-2^LEMD2/3^ "plaque". Scale bar, 5 μm. Right: Quantification of the normalized GFP::LEM-2^LEMD2/3^ integrated intensity over time from anaphase I onset to interphase for the MII chromosomal set. Error bars correspond to the standard error of the mean. The orange and grey boxes indicate interkinesis and interphase, respectively. **(F)** Representative images centered on chromosomes of fixed oocytes showing the immunolocalization of LEM-2^LEMD2/3^, DNA, and Tubulin in mid anaphase I, mid and late interkinesis. Scale bar, 5 μm.

In the earlier mid-anaphase I oocyte, vesicular membranous structures were observed on the surface of both chromosome sets. These structures formed two discontinuous double membrane layers that covered the outer surfaces. These double membrane layers were seen surrounding both the extruded chromosomal set (facing the plasma membrane of the future polar body, [PB]) and the meiosis II chromosomal set (facing the oocyte cytoplasm, [MII]). Double membrane layers were excluded from the chromosomal surfaces facing the central spindle region of both chromosomal sets. As meiosis I progressed into mid-interkinesis, the double membrane layer enveloping the MII chromosomal set displayed increased continuity, with both the mean and total lengths of membrane contours continuously expanding until late-interkinesis (Fig. 1 C, D). After extrusion of the first polar body between mid- and late-interkinesis, the mean length of membrane contours stagnated. The initial increase in mean contour length suggests that double membrane fragments likely expanded or fused to generate longer fragments. In contrast, after this initial phase, the stagnation in mean contour length, coupled with the ever-increasing total length of contours, suggested that additional membrane fragments are likely recruited to the surface of chromosomal sets between mid- and late-interkinesis. The overall length of membrane contours on the PB chromosomal set remained comparatively stable during anaphase I/interkinesis, but completely disappeared as late-interkinesis ensued (Fig. 1 C). Importantly, upon reexamination of the tomographic electron microscopy data that we had previously conducted to analyze microtubule organization during anaphase/telophase I in high-pressure frozen *C. elegans* oocytes (Laband et al., 2017), we identified identical vesicular and membranous structures on the surface of chromosomes (Fig. S1 A). The overall characteristics and dynamics of membranes were thus not significantly disrupted by the chemical fixation procedure employed in our SBF-SEM observations. These results suggest that an asymmetric double membrane structure, reminiscent of the nuclear envelope, appears progressively during interkinesis at the chromosomal surface and disappears shortly after PB extrusion.

We next probed the nature and precise kinetics of assembly of this membranous structure by time lapse imaging fertilized oocytes from a *C. elegans* transgenic strain co-expressing the nuclear envelope marker and INM LEM-domain protein LEM-2^LEMD2/3^ fused to GFP and histone H2B fused to mCherry (Fig. 1 E and Video 2) (Brachner et al., 2005; Lee et al., 2000; Lin et al., 2000). Surprisingly, LEM- 2^LEMD2/3^ localized asymmetrically to the different chromosomal sets. Unlike the double membrane structures observed in our EM analysis, LEM-2^LEMD2/3^ was only faintly detectable on the surface of the PB chromosomal set during both meiotic divisions. In stark contrast, following the onset of anaphase I, LEM-2^LEMD2/3^ gradually accumulated on the exterior surface of the MII chromosomal set, which correlated with the location of the membranous structure identified by SBF-SEM (Penfield et al., 2020). At mid- interkinesis, LEM-2^LEMD2/3^ enveloped the surface of the MII chromosomal set and reached its peak intensity. In late-interkinesis, it gradually diminished from the MII chromosomal surface, only to reappear during the onset of anaphase II. Consistent with an earlier observation, we noted a robust accumulation of LEM-2^LEMD2/3^ at the end of anaphase II on the inner (central spindle-facing) surface of the decondensing maternal pronucleus (Penfield et al., 2020). This accumulating LEM-2^LEMD2/3^ appeared to form a distinct "plaque"-like structure, which is the recruitment site of ESCRT-III complex proteins, such as CHMP-7^CHMP7^ and VPS-32^CHMP4^ (Fig. 1 E, cyan arrows) (Gatta and Carlton, 2019; Gu et al., 2017; Penfield et al., 2020). These ESCRT-III proteins are involved in the remodeling and sealing of the nuclear envelope proximal to the central spindle (Barger et al., 2023; Penfield et al., 2020). We did not observe any LEM-2^LEMD2/3^ "plaque"-like structure nor the accumulation of CHMP-7^CHMP7^ or VPS-32^CHMP4^ during interkinesis (Fig. S1 B). This observation aligned with the absence of double membrane sealing in our late interkinesis SBF-SEM reconstruction, indicating that, unlike a typical nuclear envelope, double membranes never fully enclose the MII chromosomal set during interkinesis. Therefore, we chose the term "interkinetic envelope" to describe this unique, asymmetric, and non-canonical membranous structure.

### Microtubules and proximity to the plasma membrane negatively regulate interkinetic envelope assembly

Next, we investigated the origin of the asymmetric assembly of the interkinetic envelope, which initiated on the external surface of chromosomes, and was more pronounced on the MII compared to the PB chromosomal set. We have previously demonstrated that in *C. elegans* oocytes, after anaphase onset, meiotic spindle pole microtubules disassemble before central spindle microtubules (Laband et al., 2017). This temporal uncoupling mirrors the observed asymmetry in interkinetic envelope assembly, which begins on the external (spindle pole-facing) surface of chromosomes before progressing toward the internal (central spindle-facing) surface (Figure 1 F). This observation suggested a potential functional link between microtubule disassembly and interkinetic envelope assembly, similar to nuclear envelope reformation during mitotic exit (Dey and Baum, 2021). To directly test this hypothesis, we treated oocytes with a low dose of colchicine immediately after anaphase onset to promote microtubule disassembly while allowing chromosome segregation to continue. We then monitored the recruitment of GFP-tagged LEM- 2^LEMD2/3^ as a marker for interkinetic envelope assembly (Fig. S1 C). In colchicine-treated oocytes, LEM-2^LEMD2/3^ was recruited more rapidly to the chromosome surface and formed a more continuous layer around the MII chromosomal set compared to controls. These results indicate that spindle microtubule disassembly triggers interkinetic envelope formation, and prevents premature assembly on the internal surface of chromosomes.

The visible asymmetry between the MII and PB interkinetic envelopes mirrored the uneven positioning of the two chromosomal sets during anaphase and interkinesis. Specifically, the PB chromosomes were oriented toward the plasma membrane, while the MII chromosomes faced the oocyte cytoplasm (Fig. 1 B-E). This led us to hypothesize that the plasma membrane might act as a barrier, inhibiting or delaying interkinetic envelope assembly around the PB chromosomal set. To test this, we depleted the dynein adaptor protein LIN-5^NuMA^ via RNAi (Fig. S1 D). LIN-5^NuMA^ is crucial for recruiting dynein to meiotic spindle poles, which in turn is essential for microtubule focusing at the spindle poles and proper spindle rotation perpendicular to the plasma membrane before anaphase (van der Voet et al., 2009). In LIN-5^NuMA^- depleted oocytes, the spindle remained parallel to the oocyte cortex, and chromosome segregation occurred parallel to the plasma membrane. Strikingly, this was accompanied by a symmetrization of GFP-tagged LEM-2^LEMD2/3^ levels between the two chromosomal sets compared to control oocytes, suggesting that proximity to the plasma membrane might hinder or delay LEM-2^LEMD2/3^ recruitment and interkinetic envelope assembly on the PB chromosomal set. Taken together, our results suggest that both the anaphase I central spindle microtubules and the proximity of the plasma membrane negatively regulate LEM-2^LEMD2/3^ recruitment, and likely also the formation of the interkinetic envelope.

### The interkinetic envelope contains INM but lacks ONM proteins

To determine the protein composition of the interkinetic envelope, we analyzed the localization of GFP-tagged INM and ONM proteins including: the unique *C. elegans* B-type lamin protein LMN-1^Lamin^ ^B1^, which lines the inner side of the INM (Liu et al., 2000), the second LEM-domain INM protein EMR-1^Emerin^ (Gruenbaum et al., 2002; Manilal et al., 1996; Nagano et al., 1996), the chromatin-binding protein BAF- 1^BAF^ (Barrier of Autointegration Factor) responsible for LEM-domain protein recruitment to the INM (Gorjánácz et al., 2007; Shumaker et al., 2001), the two LINC complex components, SUN-1^SUN1^ at the INM, and the KASH domain protein ZYG-12 at the ONM (Malone et al., 2003; Ungricht and Kutay, 2017), the ER signal peptidase and ONM marker SP12 (Poteryaev et al., 2005; Rolls et al., 2002), and the Ribosome-Associated Membrane Protein 4 RAMP4 (also known as Stress-associated Endoplasmic Reticulum Protein 1 or SERP1, (Lee et al., 2016) (Fig. 2 A, B, Video 3). In addition to the single lamin LMN-1^Lamin^ ^B1^, all INM proteins tested, including LEM-2^LEMD2/3^, EMR-1^Emerin^, BAF-1^BAF^, and SUN-1^SUN1^ were located at the surface of the MII chromosomal set during interkinesis colocalizing with the interkinetic envelope, with BAF-1^BAF^ also localized all over the chromosome mass. Instead, ONM proteins ZYG-12, SP12 and RAMP4 were absent (Fig. 2 A-C). In the nuclear envelope, the ONM is continuous and functionally interrelated with the ER, with which it shares numerous proteins and markers (Whaley et al., 1960). The lack of ONM markers in the interkinetic envelope suggests that, unlike canonical nuclear envelopes, the interkinetic envelope is not contiguous with the ER. We confirmed this hypothesis by analyzing the ultrastructure of the ER, close to the interkinetic envelope, using SBF-SEM (Fig. 2 D and Video 4). Although ER membrane sheets were present near the MII chromosomal set throughout anaphase and interkinesis, they were visibly distinct and physically separated from the interkinetic envelope. Thus, the interkinetic envelope on the MII chromosomal set contains INM proteins but lacks ONM proteins, likely due to its physical disconnection from the ER.

**Figure 2:**
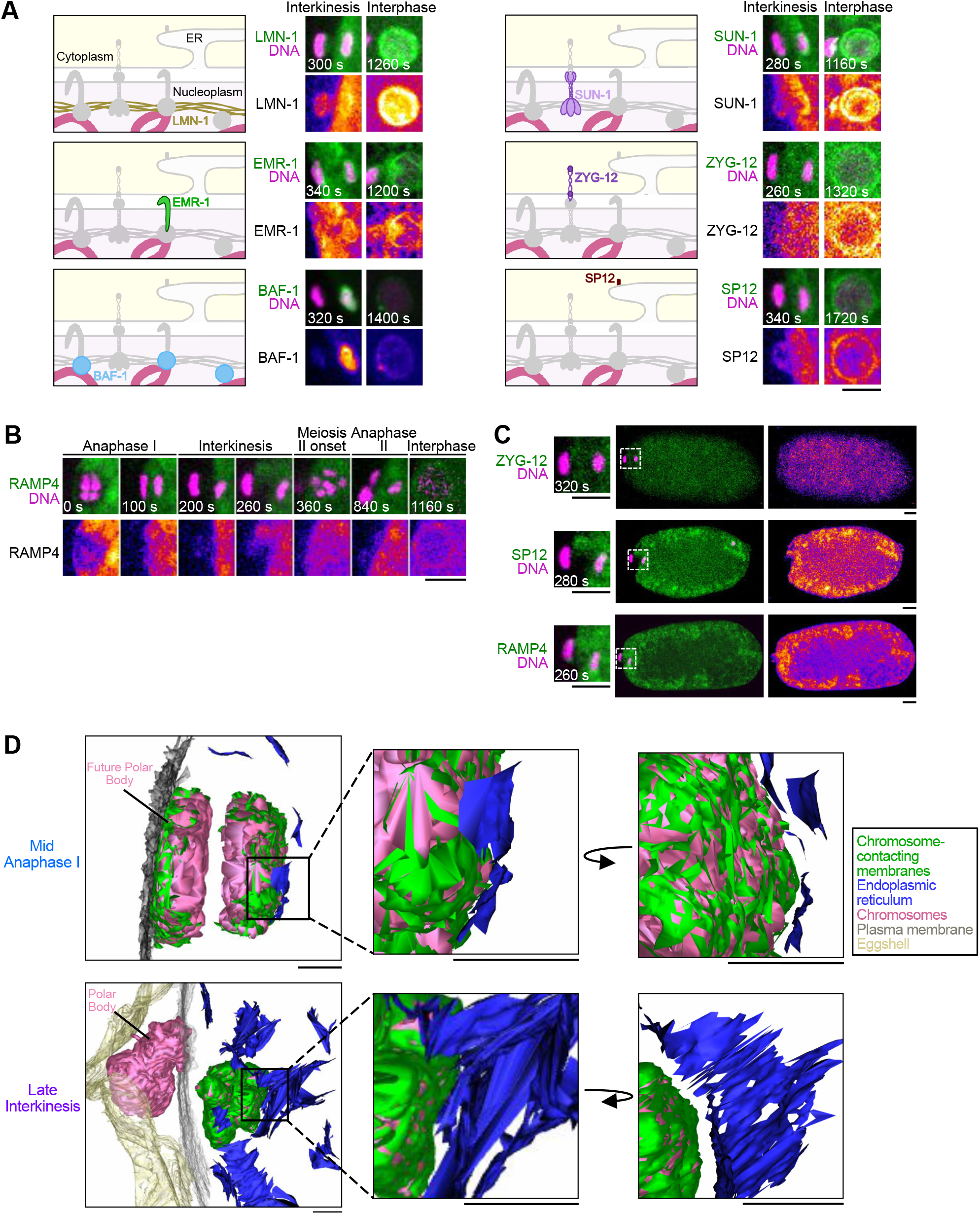
The interkinetic envelope contains INM but lacks ONM proteins. **(A)** Left: Schematics of INM and ONM protein theoretical localization at the nuclear envelope. Right: Representative images of a ROI centered around chromosomes from oocytes expressing mCherry::H2B and either GFP::EMR-1^Emerin^ (n= 6), GFP::BAF-1^BAF^ (n= 20), GFP::LMN-1^LaminB1^ (n= 7), SUN-1^SUN1^::GFP (n= 17), GFP::ZYG-12 (n= 14), or GFP::SP12 (n= 18) during interkinesis and interphase. Timings indicated at the bottom left corner of images are from anaphase I onset. Scale bar, 5 μm. **(B)** Representative time-lapse images centered on chromosomes of oocytes expressing mCherry::H2B (magenta) and GFP::RAMP4 (green) (n= 6) during meiosis I and II. Timings indicated at the bottom left corners of images are from anaphase I onset. Scale bar, 5 μm. **(C)** Representative images of oocytes expressing mCherry::H2B (magenta) and either GFP::ZYG-12, GFP::SP12 or GFP::RAMP4 (green) during interkinesis, with a magnification of the ROI (white dashed box) displayed on the left. **(D)** Left: 3-dimensional reconstructions centered on chromosomes of a mid-anaphase I (top) and a late-interkinesis (bottom) oocyte acquired by SBF-SEM. Chromosomes in magenta, membranes in contact with chromosomes in green, plasma membrane in gray, eggshell in gold, and endoplasmic reticulum in blue. Right: Magnifications of an ROI viewed from two different angles to show the lack of continuity between the interkinetic envelope and the ER. Scale bars, 1 μm for the full view and 0.5 μm for the ROI.

### BAF-1^BAF^ and VRK-1^VRK1^ control the structural integrity of the interkinetic envelope

Upon mitotic exit in both *C. elegans* and human tissue cultured cells, BAF plays a crucial role in nuclear envelope reformation (Asencio et al., 2012; Gorjánácz et al., 2007; Liu et al., 2003; Samwer et al., 2017; Schellhaus et al., 2016). To explore the potential involvement of the equivalent *C. elegans* protein in interkinetic envelope formation, we performed SBF-SEM following the full depletion of BAF-1^BAF^ in oocytes. Due to the inherent challenge of achieving full depletion through RNAi alone of a small 90 amino-acid protein such as BAF-1^BAF^, we employed a dual approach, combining RNAi with Auxin treatment in a transgenic strain engineered to express endogenous BAF-1^BAF^ fused to an AID (Auxin-Inducible Degron) tag (Zhang et al., 2015) (Fig. S2 A, B). Complete depletion of BAF-1^BAF^, achieved only through the combination of RNAi and Auxin treatments, did not inhibit the formation of the interkinetic envelope (Fig. 3 A, Fig. S2 A-C, and Video 5). However, while control oocytes displayed a nearly continuous envelope covering the outer surface of the MII chromosomal set, BAF-1^BAF^-depleted oocytes assembled a highly fenestrated envelope with a strong reduction in overall membrane density (total membrane length in contact with chromosomes in control oocytes was 325.24 µm vs. 183.82 µm in the absence of BAF-1^BAF^). Unlike in controls, in BAF-1^BAF^-depleted oocytes, small membrane fragments covered the surface of both chromosomal sets and were found inside the chromatin masses of the segregating chromosomes (Fig. 3 A, white arrows). Thus, BAF-1^BAF^ depletion leads to a drastic reduction in the recruitment of membranes necessary for interkinetic envelope assembly, coupled with strong defects in membrane fusion and distribution over both segregating chromosomal sets.

**Figure 3:**
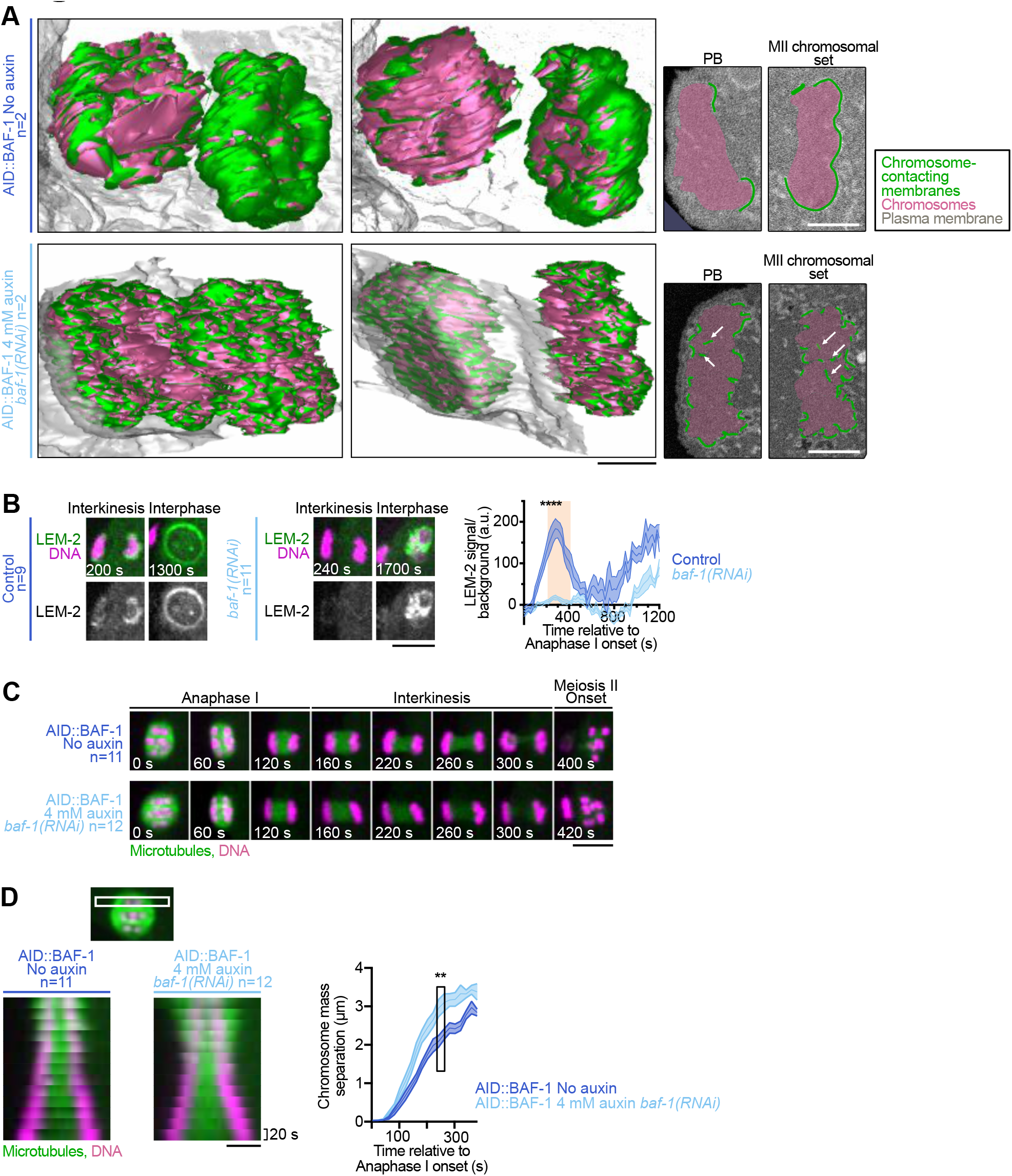
BAF-1^BAF^ is essential for the integrity of the interkinetic envelope, for INM protein localization, and for normal chromosome segregation. **(A)** Left: 3- dimensional reconstructions centered on chromosomes of a mid-interkinesis control oocyte (AID::BAF-1^BAF^, No auxin) (top) and a mid-interkinesis BAF-1^BAF^-depleted oocyte (AID::BAF-1^BAF^, 4 mM auxin, *baf-1(RNAi)*) (bottom) viewed from two different angles. Scale bar, 1 µm. Right: 2-dimensional single sections of two ROIs centered on each chromosomal set. Chromosomes in magenta, membranes in contact with chromosomes in green, and plasma membrane in gray. White arrows indicate membrane fragments within the chromosomal sets in absence of BAF-1^BAF^. Scale bar, 1 µm. **(B)** Left: Representative time-lapse images centered on chromosomes of oocytes expressing mCherry::H2B (magenta) and GFP::LEM-2^LEMD2/3^ (green) during interkinesis and interphase in the indicated conditions. Timings indicated at the bottom left corners of images are from anaphase I onset. Scale bar, 5 μm. Right: Quantification of the normalized GFP::LEM-2^LEMD2/3^ integrated intensity over time from anaphase I onset to interphase for the MII chromosomal set. Control in dark blue, *baf-1(RNAi)* in light blue. Error bars correspond to the standard error of the mean. The orange box indicates interkinesis. Mann-Whitney test on the mean value of GFP::LEM-2^LEMD2/3^ intensity in interkinesis (**** p <0.0001). **(C)** Representative time-lapse images centered on chromosomes of oocytes expressing GFP::TBA-2^α-tubulin^ (green) and mCherry::HIS-11^H2B^ (magenta) during meiosis I in the indicated conditions. Timings indicated at the bottom left corners of images are from anaphase I onset. Scale bar, 5 μm. **(D)** Left: Kymographs showing a pair of segregating chromosomes in GFP::TBA-2^α-tubulin^ (green) and mCherry::HIS-11^H2B^ (magenta)-expressing oocytes from anaphase I onset in the indicated conditions. Scale bar, 5µm. Right: Quantification of the distance between the two sets of segregating chromosomes over time from anaphase I onset in control oocytes (dark blue) and BAF-1^BAF^-depleted oocytes (light blue). Error bars correspond to the standard error of the mean. Mann-Whitney test on the mean distance between the segregating chromosomal sets during interkinesis in both conditions (** p <0.01).

During nuclear envelope reformation, BAF mediates the physical interaction between chromatin and LEM-domain proteins, which is essential for nuclear envelope integrity (Gorjánácz et al., 2007; Liu et al., 2003). To determine if the phenotype we observed upon BAF-1^BAF^ depletion could be attributed to defects in LEM-domain protein recruitment, we employed RNAi to knock down BAF-1^BAF^ and monitored the presence of GFP-tagged INM proteins LEM-2^LEMD2/3^ and EMR-1^Emerin^ (Fig. 3 B, Fig. S2 D, E and Video 6). Depletion of BAF-1^BAF^ resulted in a significant reduction of both INM proteins from the interkinetic envelope. Together, these results suggest that BAF-1^BAF^ plays an important role in controlling the integrity and continuity of the interkinetic envelope, potentially by recruiting LEM-domain proteins on the chromatin surface. The structural integrity defects observed in the interkinetic envelope following BAF-1^BAF^ depletion prompted us to examine its potential impact on chromosome segregation. For this, we conducted time lapse imaging of oocytes expressing GFP-tagged tubulin and mCherry-tagged H2B, with and without BAF-1^BAF^ (Fig. 3 C). In both conditions, chromosomes aligned at the spindle equator on a tight metaphase plate during metaphase I. Throughout anaphase I, the segregating chromosomes maintained a compact arrangement, showing no signs of mis-segregation in both control and BAF-1^BAF^-depleted oocytes. On the other hand, chromosome segregation was noticeably faster and resulted in a significantly increased distance between the two segregating chromosomal sets in the absence of BAF-1^BAF^ compared to control oocytes (Fig. 3 D). Since the overall chromosome structure and condensation, and the meiosis I spindle organization appeared normal in the absence of BAF-1 (Fig. S2 F, G), our results suggest that the integrity of the interkinetic envelope impacts the normal pace and extent of chromosome segregation in *C. elegans* oocytes.

Since the complete depletion of BAF-1^BAF^ resulted in the formation of a highly fenestrated interkinetic envelope with significantly reduced membrane content, we next investigated whether over-recruiting BAF-1^BAF^ on chromosomes would have the opposite effect. To test this, we depleted the VRK-1^VRK1^ kinase, which phosphorylates BAF-1^BAF^ at mitotic entry to promote its detachment from chromatin—an event essential for efficient nuclear envelope breakdown (NEBD) (Fig. 4 A)(Gorjánácz et al., 2007). In the absence of VRK-1^VRK1^ during mitosis, BAF-1^BAF^ remains permanently bound to chromatin, leading to defects in NEBD and nuclear envelope reformation after mitosis, with excess membranes forming around chromosomes(Asencio et al., 2012; Gorjánácz et al., 2007). First, we verified the presence of VRK-1^VRK1^ at the surface of oocyte chromosomes during interkinesis (Fig. 4 B). Then, we confirmed that, similar to mitosis, depleting VRK-1^VRK1^ led to an excess of BAF-1^BAF^ and LEM-2^LEMD2/3^ on chromosomes during interkinesis (Fig. 4 C, D). Notably, in the absence of VRK-1^VRK1^, GFP-tagged BAF-1^BAF^ and LEM-2^LEMD2/3^ were strongly recruited to both the PB and MII chromosomal sets, unlike in control oocytes. This suggests that VRK-1 is at least partially responsible for the asymmetric localization of both proteins on chromosomes during interkinesis. Moreover, in the absence of VRK-1^VRK1^, this overaccumulation of BAF-1^BAF^ and LEM-2^LEMD2/3^ on chromosomes was accompanied by their noticeable stretching during segregation, which is reminiscent of the phenotype observed during mitosis in the same condition (Fig. 4 E). These defects of mitotic chromosome segregation have previously been attributed to the excess of membranes that surrounds them in absence of VRK-1^VRK-1^ (Gorjánácz et al., 2007). To investigate whether membrane hyper-recruitment was responsible for chromosome stretching during interkinesis in VRK-1^VRK1^-depleted oocytes, we conducted SBF-SEM followed by 3D reconstruction (Fig. 4 F). Surprisingly, although this approach confirmed the stretched chromosome phenotype, the density of membranes at the chromosomal surface was significantly reduced. In control oocytes, the total membrane length in contact with chromosomes was 325.24 µm, compared to 199.51 µm in the absence of VRK-1^VRK-1^. Thus, the chromosomal stretching observed in the absence of VRK-1^VRK1^ is likely due to a function separate from its role in interkinetic envelope assembly. Furthermore, the interkinetic envelope appeared highly fenestrated, similar to the phenotype observed in BAF-1^BAF^-depleted oocytes. Thus, our results demonstrate that both depletion and over-recruitment of BAF-1 on chromosomes lead to a similar reduction in membrane density and a highly fenestrated appearance at the chromosome surface during interkinesis. Overall, these findings underscore the critical role of BAF-1, whose chromosomal levels must be tightly regulated to ensure the proper assembly of the interkinetic envelope.

**Figure 4:**
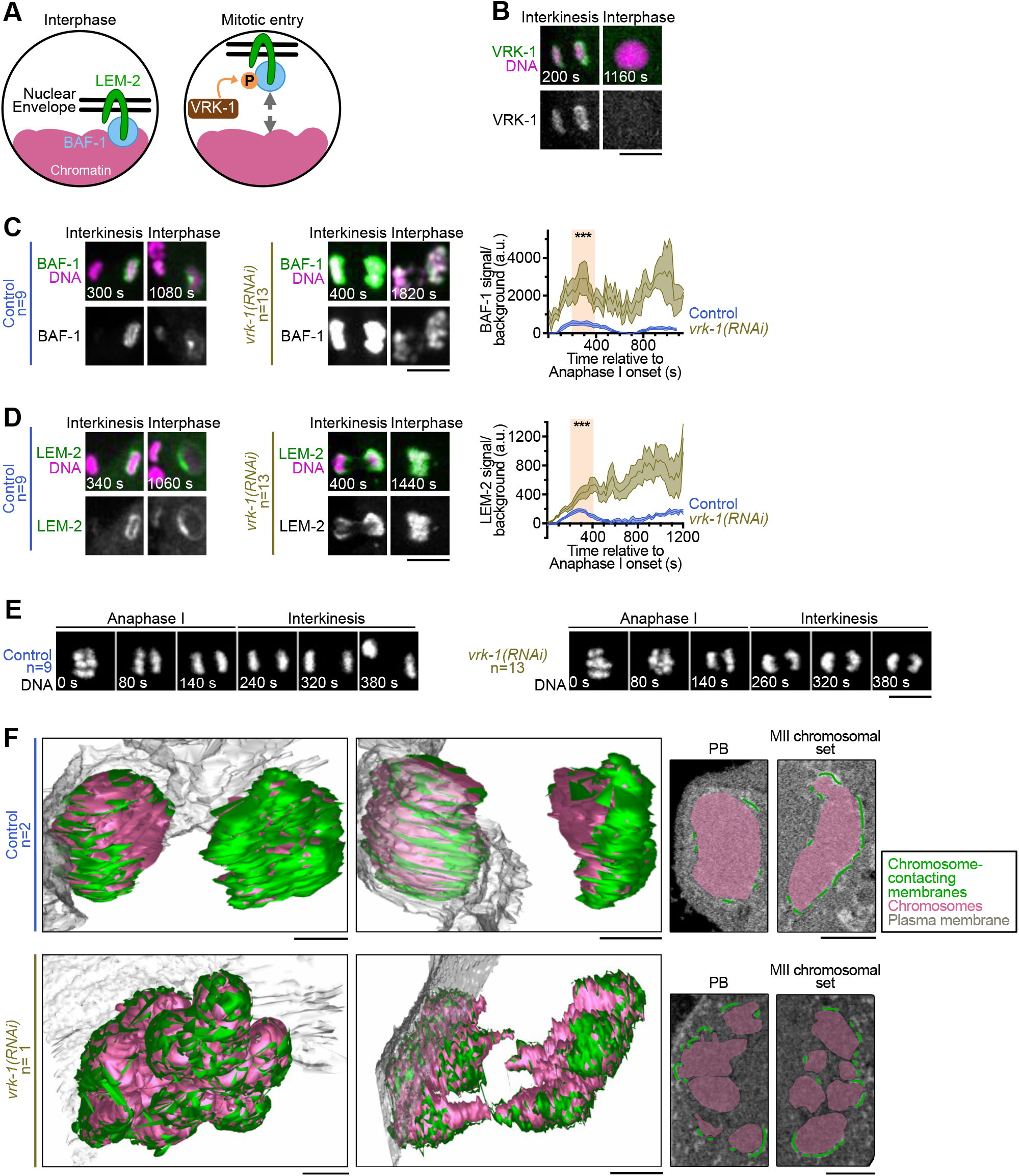
VRK-1^VRK^ is essential for regulating BAF-1^BAF^ and LEM-2^LEMD2/3^ recruitment and for the integrity of the interkinetic envelope. **(A)** Left: Schematics of BAF-1^BAF^ and LEM-2^LEMD2/3^ localization in interphase. Right: At mitotic entry VRK-1^VRK1^ phosphorylates BAF-1^BAF^ and disrupts its chromatin binding. **(B)** Representative images of a ROI centered around chromosomes from oocytes expressing GFP::H2B and VRK-1^VRK1^::mCherry (n= 12) during interkinesis and interphase. Timings indicated at the bottom left corner of images are from anaphase I onset. Scale bar, 5 μm. **(C, D)** Left: Representative time-lapse images centered on chromosomes of oocytes expressing mCherry::H2B (magenta) and (C) GFP::BAF- 1^BAF^ or (D) GFP::LEM-2^LEMD2/3^ (green) during interkinesis and interphase in the indicated conditions. Timings indicated at the bottom left corners of images are from anaphase I onset. Scale bar, 5 μm. Right: Quantification of the normalized GFP::LEM-2^LEMD2/3^ integrated intensity over time from anaphase I onset to interphase for the MII chromosomal set. Control in dark blue and *vrk-1(RNAi)* in light brown. Error bars correspond to the standard error of the mean. The orange box indicates interkinesis. Mann-Whitney test on the mean value of GFP::LEM-2^LEMD2/3^ intensity in interkinesis (*** p<0.001). **(E)** Representative time-lapse images centered on chromosomes of oocytes expressing mCherry::H2B (gray) during anaphase I and interkinesis in the indicated conditions. Timings indicated at the bottom left corners of images are from anaphase I onset. Scale bar, 5 μm. **(F)** Left: 3-dimensional reconstructions centered on chromosomes of a control oocyte (top) and a VRK-1^VRK1^- depleted oocyte (bottom) viewed from two different angles. Scale bar, 1 µm. Right: 2- dimensional single sections of two ROIs centered on each chromosomal set. Chromosomes in magenta, membranes in contact with chromosomes in green, and plasma membrane in gray. Scale bar, 1 µm.

#### MEL-28^ELYS^ is essential for membrane recruitment on the chromosomal surface

We next sought to determine the origin of the double membranes that form the interkinetic envelope. In addition to BAF, nuclear envelope reformation at the end of mitosis requires the nucleoporin ELYS (Franz et al., 2007; Galy et al., 2006; Rasala et al., 2006). ELYS is a member of the ’Y’ nuclear pore subcomplex, which is essential for post-mitotic NPC reassembly and nuclear envelope integrity (Harel et al., 2003; Walther et al., 2003). In *C. elegans*, the orthologous protein MEL-28^ELYS^ is essential for nuclear envelope formation and function in embryos, and for chromosome segregation in oocytes (Fernandez and Piano, 2006; Galy et al., 2006; Gomez-Saldivar et al., 2016; Hattersley et al., 2016). During metaphase I in *C. elegans* oocytes, MEL-28^ELYS^ colocalizes with kinetochore cup-like structures on the surface of chromosomes, where it serves as a docking site for the catalytic subunit of protein phosphatase 1 (PP1c) (Hattersley et al., 2016). In anaphase I, the localization of MEL-28^ELYS^ expanded to all chromosomes, while also maintaining colocalization with the interkinetic envelope on both chromosomal sets (Fig. 5 A). At this stage, PP1c orchestrates kinetochore disassembly, a critical process for proper chromosome segregation (Hattersley et al., 2016). To explore the potential involvement of MEL-28^ELYS^ in interkinetic envelope assembly and double membrane recruitment, we performed SBF-SEM following RNAi-mediated depletion of MEL- 28^ELYS^. In line with previous findings, in the absence of MEL-28^ELYS^, chromosomes within each chromosomal set were not as tightly grouped as compared to in controls during anaphase I (Fig. 5 B, Fig. S3 A and Video 7). Furthermore, physical segregation of the two chromosomal sets aborted rapidly after anaphase onset leading to a shorter separation distance compared to in control oocytes at the same stage. Thus, to confirm that MEL-28^ELYS^-depleted oocytes had reached interkinesis at the time of chemical fixation, we used time lapse microscopy to capture their dynamics before proceeding with fixation and SBF-SEM (Fig. 5 B, C, Fig. S3 B, C and Video 8). After segmentation and 3D reconstruction, we observed a drastic reduction in double membranes on the outer surface of chromosomes in MEL-28^ELYS^- depleted oocytes compared to in controls (total membrane length in contact with chromosomes in control oocytes was 317.53 µm vs. 17.58 µm in the absence of MEL-28^ELYS^). Upon segmentation and reconstruction of the other membrane compartments surrounding the segregating chromosomes (vesicles, mitochondria, and ER), we observed the expected meiotic spindle organelle exclusion zone around chromosomes in control oocytes (Fig. 5 D, Fig. S3 C and Video 8) (Albertson and Thomson, 1993). However, in the absence of MEL-28^ELYS^, the organelle exclusion zone was notably wider. Furthermore, this zone encompassed numerous unidentified small membrane fragments that surrounded the chromosomes but did not directly contact them—both features not observed in control oocytes. Thus, MEL-28^ELYS^ is required for interkinetic envelope assembly.

**Figure 5:**
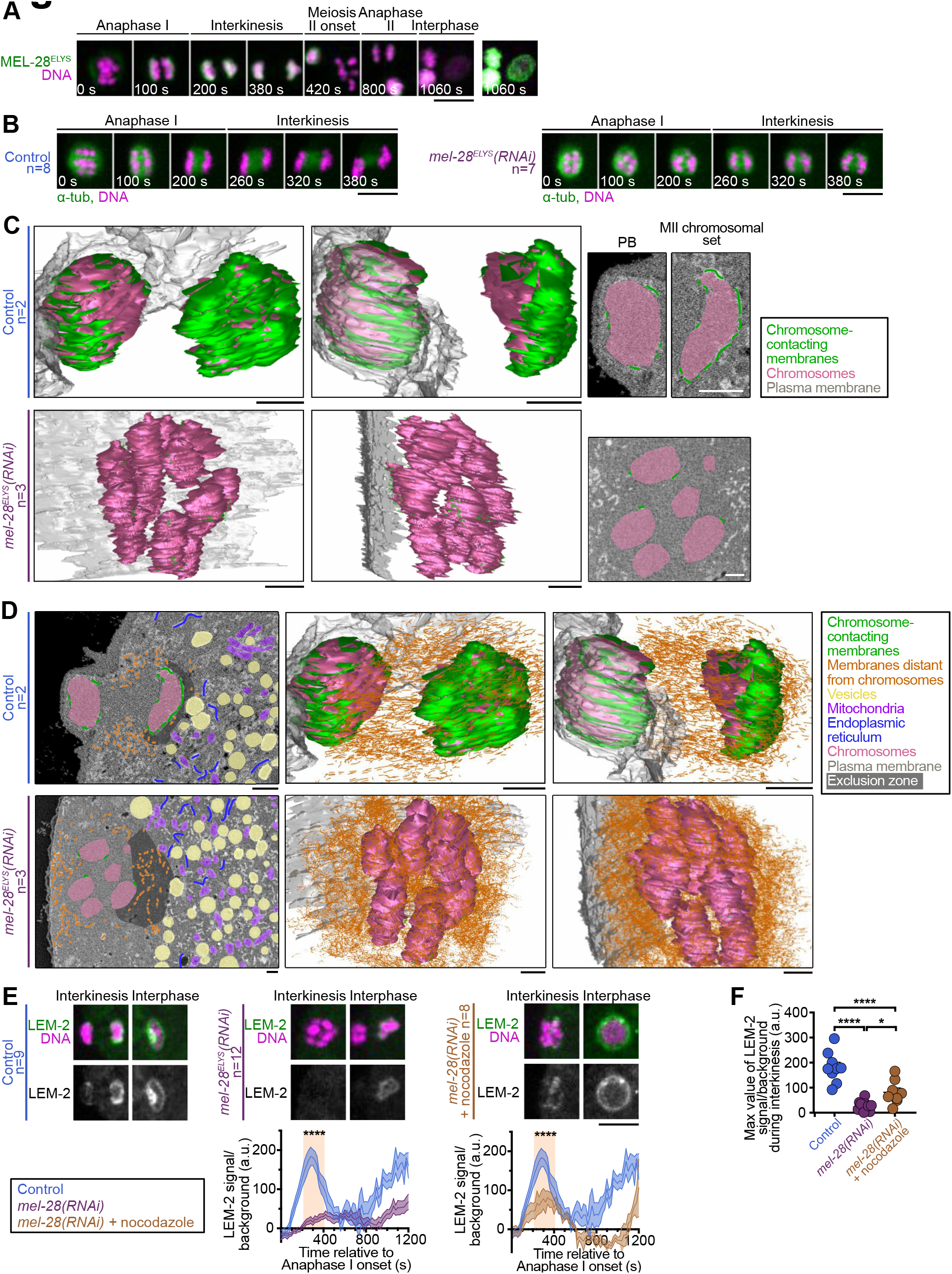
MEL-28^ELYS^ is essential for the interkinetic envelope integrity and membrane recruitment. **(A)** Representative time-lapse images centered on chromosomes of oocytes (n= 8) expressing mCherry::H2B (magenta) and GFP::MEL-28^ELYS^ (green) during meiosis I and II. Timings indicated at the bottom left corners of images are from anaphase I onset. Scale bar, 5 μm. **(B)** Representative time-lapse images centered on chromosomes of oocytes expressing mCherry::H2B (magenta) and GFP::TBA-2^α-tubulin^ (green) and mCherry::HIS-11^H2B^ (green) during anaphase I and interkinesis in the indicated conditions. Timings indicated at the bottom left corners of images are from anaphase I onset. Scale bar, 5 μm **(C)** Left: 3- dimensional reconstructions centered on chromosomes of a control oocyte (top) and a MEL-28^ELYS^-depleted oocyte (bottom) viewed from two different angles. Scale bar, 1 µm. Right: 2-dimensional single sections of ROIs centered on each chromosomal set. Chromosomes in magenta, membranes in contact with chromosomes in green, and plasma membrane in gray. Scale bar, 1 µm. **(D)** Left: 2-dimensional single sections of ROI centered on each chromosomal set of a control oocyte (top) and a MEL-28^ELYS^-depleted oocyte (bottom). Scale bar, 1 µm. Right: 3-dimensional reconstructions centered on chromosomes viewed from two different angles. Scale bar, 1 µm. Chromosomes in magenta, membranes in contact with chromosomes in green, membranes distant from chromosomes in orange, vesicles in yellow, mitochondria in purple, endoplasmic reticulum in blue, and plasma membrane in gray. Scale bar, 1 µm. **(E)** Top: Representative time-lapse images centered on chromosomes of oocytes expressing mCherry::H2B (magenta) and GFP::LEM- 2^LEMD2/3^ (green) during interkinesis and interphase in the indicated conditions. Timings indicated at the bottom left corners of images are from anaphase I onset. Scale bar, 5 μm. Bottom: Quantification of the normalized GFP::LEM-2^LEMD2/3^ integrated intensity over time from anaphase I onset to interphase for the MII chromosomal set. Control in dark blue, *mel-28(RNAi)* in purple, *mel-28(RNAi)* treated with 100 ng/µL nocodazole in brown. Error bars correspond to the standard error of the mean. The orange box indicates interkinesis. Mann-Whitney test on the mean value of GFP::LEM-2^LEMD2/3^ intensity in interkinesis (**** p <0.0001). **(F)** Quantifications of the maximal mean value of GFP::LEM-2^LEMD2/3^ intensity in interkinesis normalized over background in the indicated conditions. One-way Anova test (**** p <0.0001, * p <0,05).

We hypothesized that the expanded organelle exclusion zone and the deficiencies in interkinetic envelope assembly in the absence of MEL-28^ELYS^ might, at least in part, result from the abnormal persistence of spindle pole microtubules throughout anaphase and interkinesis (Fig. 5 B) (Hattersley et al., 2016). These ectopic spindle pole microtubules could potentially act as a physical barrier, preventing membrane recruitment on the surface of chromosomes. To test this hypothesis, we compared the intensity of GFP::LEM-2^LEMD2/3^ around chromosomes during interkinesis in the absence of MEL-28^ELYS^, with and without microtubule depolymerization induced by nocodazole treatment (Fig. 5 E, F and Fig. S3 D). As expected, GFP::LEM-2^LEMD2/3^ was nearly absent from the surface of both chromosomal sets in the absence of MEL-28^ELYS^. Importantly, this absence of GFP::LEM-2^LEMD2/3^ was not attributed to the delocalization of BAF-1^BAF^, which remained properly localized on the MII chromosomal set in the absence of MEL- 28^ELYS^ (Fig. S3 E). Furthermore, upon microtubule depolymerization induced by nocodazole, GFP::LEM-2^LEMD2/3^ levels on the MII chromosomal set were partially restored, reaching approximately half of the levels observed in control oocytes. Therefore, microtubule depolymerization can partially ameliorate the defects in interkinetic envelope assembly induced by MEL-28^ELYS^ depletion. During post-mitotic nuclear envelope reformation, ER sheets are enlisted at the chromosome surface to serve as a membrane source (Anderson and Hetzer, 2008; Anderson et al., 2009; Barger et al., 2022; Deolal et al., 2024; Haraguchi et al., 2001; Otsuka et al., 2018). Our findings collectively propose a distinct mechanism for interkinetic envelope assembly, implicating the MEL-28^ELYS^-mediated accumulation of small unidentified membrane fragments on surface of the chromosomal sets. Moreover, this process is contingent, at least to some extent, on the rapid disassembly of spindle pole microtubules, a process that occurs early during anaphase in control oocytes, but is strongly delayed in the absence of MEL-28^ELYS^ (Hattersley et al., 2016).

### The interkinetic envelope contains nucleoporins but not NPCs

MEL-28^ELYS^ is essential for post-mitotic nuclear pore complex reformation (Franz et al., 2007; Galy et al., 2006). At mitotic exit, MEL-28^ELYS^ binds to chromatin and recruits other nuclear pore components that form 6-8 protein modules, the NPC subcomplexes (Fernandez-Martinez and Rout, 2021; Huang et al., 2023; Lin and Hoelz, 2019). In *C. elegans*, the NPC comprises 28 identified nuclear pore proteins (NPPs) distributed into 6 subcomplexes: the cytoplasmic and nucleoplasmic rings (also known as the ’Y-complex’, which contains MEL-28^ELYS^), the inner ring, the transmembrane nucleoporins, the central channel, the nuclear basket, and the cytoplasmic filaments (Cohen-Fix and Askjaer, 2017). In addition to MEL-28^ELYS^, previous work revealed the presence of NPP-6^NUP160^, a Y-complex nucleoporin, on the surface of the segregating chromosomal sets during anaphase I in *C. elegans* oocytes (Penfield et al., 2020).

Using transgenic and endogenously fluorescently-tagged *C. elegans* strains, we analyzed the interkinetic localization of 18 nucleoporins (of the 28 *C. elegans* NPPs) that systematically represent all major subcomplexes of the NPC (Fig. 6). We first validated NPP-6^NUP160^ localization and observed a comparable chromosomal pattern for other Y-complex constituents, including NPP-2^NUP85^, NPP-5^NUP107^, NPP- 15^NUP133^, NPP-18^SEH1^, and NPP-20^SEC13R^. The inner ring complex lines the inner part of the NPC. We detected the presence of the inner ring component NPP-19^NUP35^, but not NPP-13^NUP93^ or NPP-8^NUP155^, at the outer surface of the MII chromosomal set. The central channel is formed by nucleoporins that contain FG repeats essential for establishing nuclear pore permeability. We could not detect the presence of central channel NPP-1^NUP54^, and NPP-11^NUP62^ predominantly localized between the two sets of segregating chromosomes in a region corresponding to the anaphase I central spindle. The nuclear basket forms the nucleoplasmic side of the NPC. We found nuclear basket NPP-7^NUP153^ distributed across the entire surface of both chromosome sets, whereas NPP-21^TPR^ was absent. On the other side of the NPC, the cytoplasmic filaments include the nucleoporin NPP-24^NUP88^, which like NPP-11^NUP62^, was concentrated in the central spindle region during interkinesis. Finally, transmembrane nucleoporins NPP-12^NUP210^ and NPP-25^TMEM33^, but not NPP-22^NDC1^, were present on the outer surface of the MII chromosomal set. In summary, our investigation revealed the presence of all examined Y-complex nucleoporins, along with specifically NPP- 19^NUP35^ (inner ring), NPP-12^NUP210^ and NPP-7^NUP153^ (cytoplasmic filaments), and NPP-25^TMEM33^ (transmembrane), distributed across the entire surface of both chromosomal sets and/or asymmetrically at the interkinetic envelope. That is, nucleoporins within the same NPC subcomplex were not necessarily co-recruited to the interkinetic envelope, preventing the formation of functional subcomplexes or NPCs. In agreement, despite the presence of various nucleoporins and in alignment with prior observations (Penfield et al., 2020), both our SBF-SEM and electron tomography analyses consistently indicated an absence of nuclear pores within the interkinetic envelope. Overall, while lacking nuclear pores, the presence of nucleoporins hints at their potential engagement in unconventional functions during interkinetic envelope assembly.

**Figure 6:**
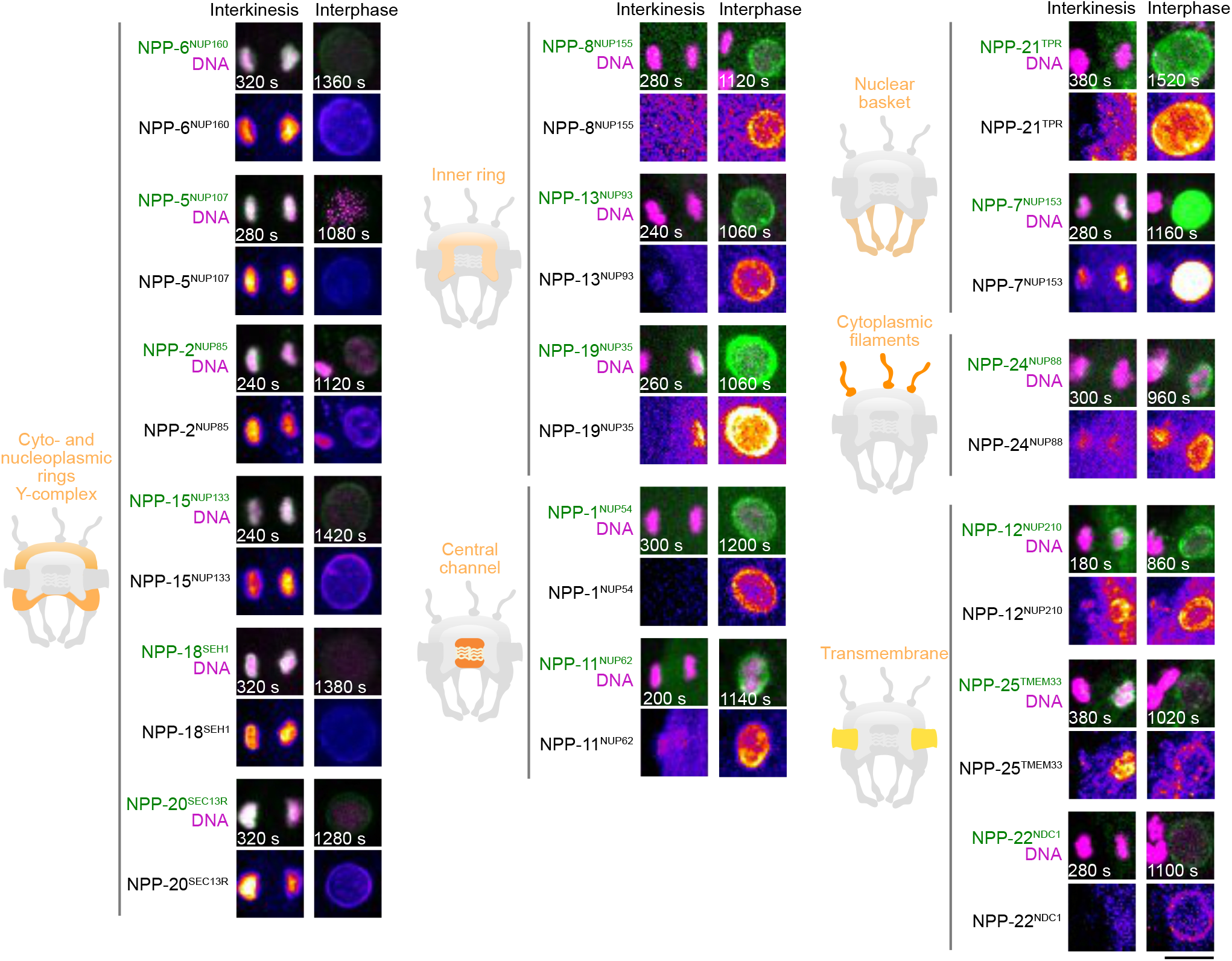
The interkinetic envelope contains nucleoporins but not NPCs. Localization of NPPs during interkinesis and interphase grouped by theoretical subcomplexes. Left: Schematics of each subcomplex localization at the NPC. Right: Representative images of a region of interest centered around chromosomes of oocytes expressing mCherry::H2B and either GFP-tagged NPP-2^NUP85^ (n= 10), NPP- 5^NUP107^ (n= 7), NPP-6^NUP160^ (n= 6), NPP-15^NUP133^ (n= 10), NPP-18^SEH1^ (n= 5), NPP-20^SEC13R^ (n= 10), NPP-13^NUP93^ (n= 6), NPP-19^NUP35^ (n= 10), NPP-12^NUP210^ (n= 7), NPP-22^NDC1^, NPP-25^TMEM33^ (n= 7), NPP-1^NUP54^ (n= 15), NPP-11^NUP62^ (n= 11), NPP- 7^NUP153^ (n= 15), NPP-21^TPR^ (n= 7), NPP-24^NUP88^ (n= 7) or mCherry-tagged NPP-8^NUP155^ (n= 14) (green) during interkinesis and interphase. Timings indicated at the bottom left corner of images are from anaphase I onset. Scale bar, 5 μm.

### Nucleoporins with membrane binding domains could contribute to interkinetic envelope integrity

We next tested the role of nucleoporins in interkinetic envelope assembly. Interestingly, aside from the two transmembrane proteins NPP-12^NUP210^ and NPP- 25^TMEM33^, responsible for post-mitotic NPC anchoring in nuclear membranes and found at the interkinetic envelope, several nucleoporins identified in the interkinetic envelope are predicted to possess domains capable of folding as amphipathic helices, which can bind to membranes (Cohen et al., 2003; Floch et al., 2015; Greber et al., 1990; Hamed and Antonin, 2021; Vollmer et al., 2015; Vollmer et al., 2012). These included Y-complex nucleoporins NPP-6^NUP160^ and NPP-15^NUP133^, inner ring component NPP-19^NUP35^, and nuclear basket protein NPP-7^NUP153^. We systematically depleted each of these six nucleoporins by RNAi and analyzed interkinetic envelope integrity by time lapse imaging using GFP::LEM-2^LEMD2/3^ intensity as a proxy (Fig. 7 A, B, S4 and Video 9). Individual depletion of all six nucleoporins led to a mild but significant decrease in GFP::LEM-2^LEMD2/3^ intensity at the chromosomal surface during interkinesis. Importantly, depleting the inner ring nucleoporin NPP-8^NUP155^, which we did not find localized at the interkinetic envelope, exhibited no discernible effect on GFP::LEM-2^LEMD2/3^ intensity (Fig. S5 A). In line with the mild decrease in LEM-2^LEMD2/3^ intensity, none of the individual nucleoporin depletions caused chromosome segregation defects (Fig. S5 B). To determine whether depleting multiple nucleoporins would have a stronger effect, we systematically co-depleted the six nucleoporins in pairs. While all co-depletions consistently exacerbated the delocalization of GFP::LEM-2^LEMD2/3^ from the chromosome surface compared to single depletions, none resulted in its complete absence (Fig. 7 B, Fig. S5 C). We were unable to assess the effect of the simultaneous depletion of NPP-12^NUP210^ and NPP-19^NUP35^, as it caused the failure of oocyte NEBD and blocked meiotic divisions. Overall, our results suggest that nucleoporins function in parallel for membrane recruitment, and their roles in interkinetic envelope assembly are at least partially redundant.

**Figure 7:**
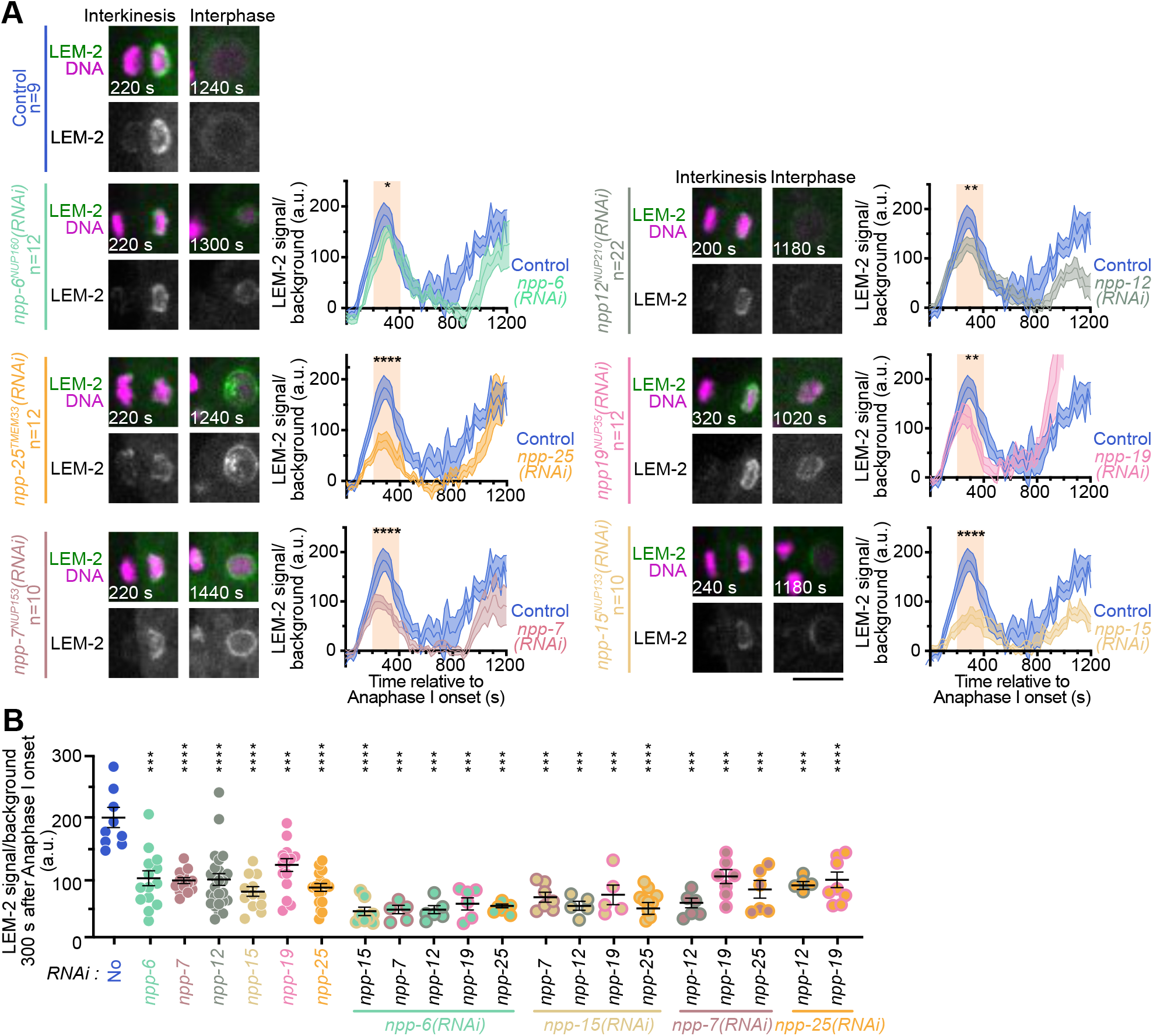
Nucleoporins with a membrane-binding domain could contribute to interkinetic envelope integrity. **(A)** Left: Representative time-lapse images centered on chromosomes of oocytes expressing mCherry::H2B (magenta) and GFP::LEM-2^LEMD2/3^ (green) during interkinesis and interphase in the indicated conditions. Timings indicated at the bottom left corners of images are from anaphase I onset. Scale bar, 5 μm. Right: Quantification of the normalized GFP::LEM-2^LEMD2/3^ integrated intensity over time from anaphase I onset to interphase for the MII chromosomal set. Control in dark blue, *npp-6^NUP160^(RNAi)* in turquoise*, npp- 25^TMEM33^(RNAi)* in dark orange, *npp-15^NUP133^(RNAi)* in light orange, *npp-12^NUP210^(RNAi)* in gray, *npp-19^NUP53^(RNAi)* in pink and *npp-7^NUP153^ (RNAi)* in light brown. Error bars correspond to the standard error of the mean. The orange box indicates interkinesis. Mann-Whitney test on the mean value of GFP::LEM-2^LEMD2/3^ intensity in interkinesis (* p <0.05, ** p<0.01, **** p<0.0001). **(B)** Quantification of the normalized GFP::LEM-2^LEMD2/3^ intensity at 300 s after anaphase I onset in the indicated conditions. Mann-Whitney test (*** p<0.001 and **** p<0.0001).

Finally, we tested whether this network of nucleoporins was hierarchically positioned downstream of MEL-28^ELYS^, akin to during post-mitotic NPC reformation (Franz et al., 2007; Galy et al., 2006). For this, we analyzed the localizations of GFP-tagged NPP-6^NUP160^, NPP-15^NUP133^, NPP-25^TMEM33^, NPP-12^NUP210^, NPP-19^NUP35^ and NPP-7^NUP153^ in oocytes during interkinesis upon MEL-28^ELYS^ depletion by RNAi (Fig. 8 and Video 10). The intensities of all six nucleoporins on the chromosome surface were markedly reduced in the absence of MEL-28^ELYS^ during interkinesis, with NPP- 6^NUP160^, NPP-7^NUP153^, and NPP-19^NUP35^ absent from chromosomes. These results indicated that chromatin-bound MEL-28^ELYS^ serves as a precursor to a network of nucleoporins including NPP-6^NUP160^, NPP-7^NUP153^, NPP-12^NUP210^, NPP-15^NUP133^, NPP- 19^NUP35^, and NPP-25^TMEM33^. This network could interact with membranes and mediate their recruitment to the chromosomal surface, thus promoting interkinetic envelope assembly.

**Figure 8:**
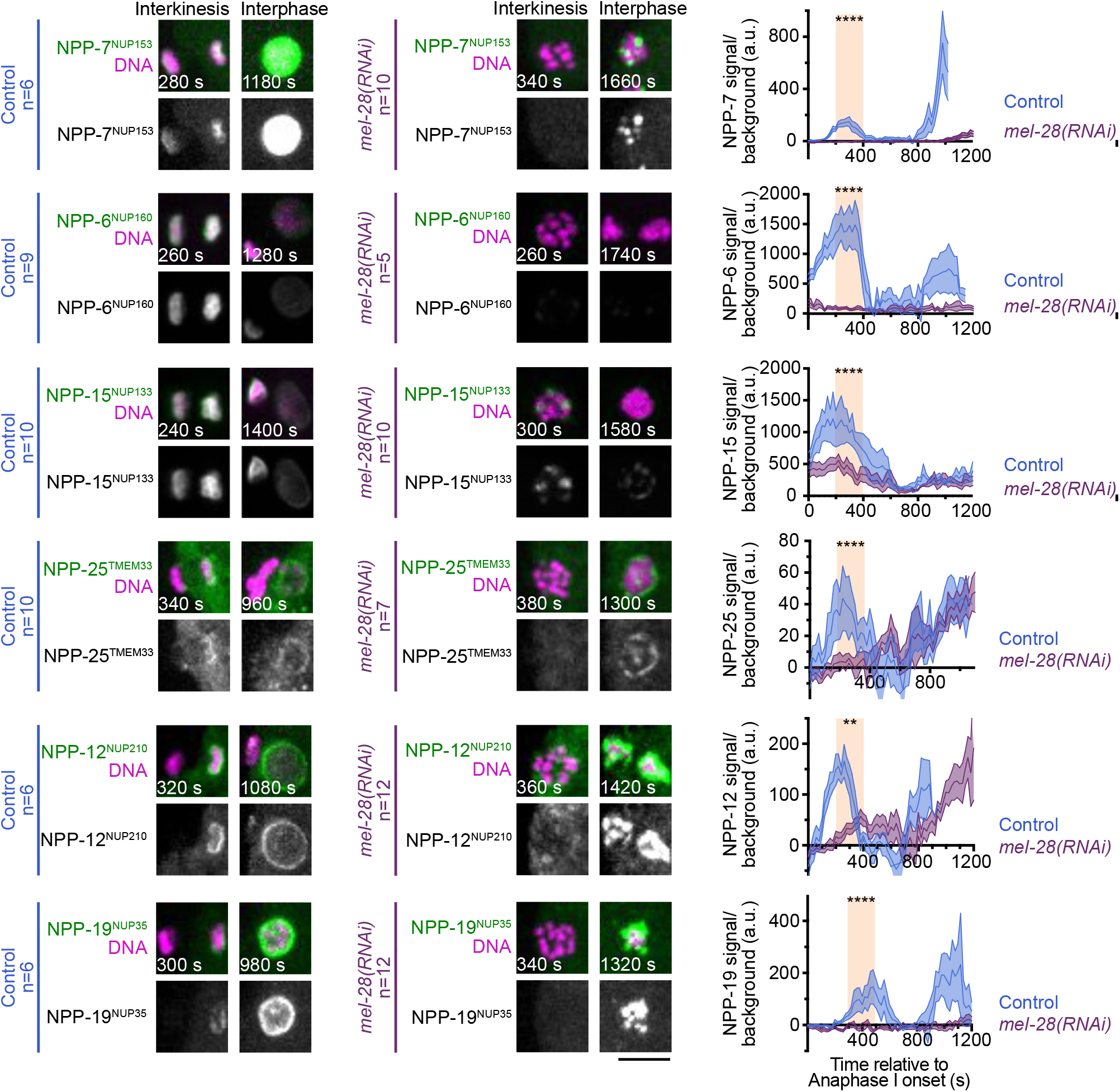
Hierarchical relationships between MEL-28^ELYS^ and nucleoporins bearing a membrane-binding domain during interkinetic envelope assembly. Left: Representative time-lapse images centered on chromosomes of oocytes expressing mCherry::H2B (magenta) and either GFP-tagged NPP-6^NUP160^, NPP- 15^NUP133^, NPP-25^TMEM33^, NPP-12^NUP210^, NPP-19^NUP35^ or NPP-7^NUP153^ (green) during interkinesis and interphase in the indicated conditions. Timings indicated at the bottom left corners of images are from anaphase I onset. Scale bar, 5 μm. Right: Quantification of the normalized GFP-tagged NPP-6^NUP160^, NPP-15^NUP133^, NPP- 25^TMEM33^, NPP-12^NUP210^, NPP-19^NUP35^ or NPP-7^NUP153^ integrated intensity over time from anaphase I onset to interphase for the MII chromosomal set. Control in dark blue and *mel-28(RNAi)* in purple. Error bars correspond to the standard error of the mean. The orange box indicates interkinesis. Mann-Whitney test on the mean value of GFP::LEM-2^LEMD2/3^ intensity in interkinesis (** p<0.01, *** p<0.001, **** p<0.0001).

## DISCUSSION

The suppression of most interphasic events during interkinesis in oocytes, including chromosome decondensation and genome replication, led to the widely accepted assumption that the nuclear envelope does not reassemble during this short transition phase between meiosis I and II (Gerhart et al., 1984; Lenart and Ellenberg, 2003; Nakajo et al., 2000; Nebreda and Ferby, 2000). We show here that, although not erroneous, this prediction is overstated. By combining electron microscopy and time lapse imaging in *C. elegans* oocytes during interkinesis, we found that an ’interkinetic envelope’ transiently forms around condensed chromosomes at this stage. Although this envelope is not nuclear as it does not compartmentalize the genome, it nevertheless shares several features with the nuclear envelope, including its double-membrane structure and protein composition of the inner layer. It also displays distinct and surprising differences from nuclear envelopes.

A striking feature of the interkinetic envelope is its lack of continuity with the ER. During post-mitotic nuclear envelope reformation, ER membranes are recruited at chromosome surfaces to regenerate nuclear envelope membranes (Anderson and Hetzer, 2008; Anderson et al., 2009; Deolal et al., 2024; Haraguchi et al., 2001; Otsuka et al., 2018). Then, because the ER membrane is contiguous with the nuclear envelope, proteins can translocate seamlessly from the ER to the ONM, resulting in partial sharing of protein composition between these structures (Deolal et al., 2024). In contrast, the lack of continuity between the interkinetic envelope and the meiotic ER likely explains the apparent absence of ONM protein within the interkinetic envelope. The reasons for the absence of a junction between the interkinetic envelope and the ER remain unclear. We propose two hypotheses to explain this lack of continuity. First, there may be an unknown physical barrier that prevents the interkinetic envelope from incorporating ER-derived membranes. Alternatively, missing components within the interkinetic envelope could inhibit the fusion of these two structures. Notably, despite being discovered decades ago, the mechanism for junction formation between the ER and the nuclear envelope in mitotic cells remains unknown (Watson, 1955; Whaley et al., 1960). Further investigation will be required to uncover the molecular mechanism underlying this unique feature of the interkinetic envelope.

Nevertheless, we demonstrated that the interkinetic envelope initially assembles on the external surface of the MII chromosomal set that will persist in the oocyte cytoplasm for meiosis II, before almost completely covering the surface of this chromosomal set. In the absence of physical contact with the ER, we found that a population of small membrane fragments, the origin and identity of which are at present unclear, positioned near chromosomes, seemed to participate in interkinetic envelope assembly. We suspect that these small membrane fragments could originate from nuclear envelope remnants following NEBD of the diakinesis oocyte (Lenart and Ellenberg, 2003). Our functional analysis suggests that MEL-28^ELYS^ acts as an upstream regulator of these fragments. In its absence, the small membrane fragments concentrated around the segregating chromosomes but did not contact them to assemble an envelope at their surface.

We identified two complementary functions for MEL-28^ELYS^ at the interkinetic envelope. First, we found that in absence of MEL-28^ELYS^, the observed small membrane fragments were positioned further away from chromosomes, likely caused by persistent ectopic spindle poles during anaphase I in absence of MEL-28^ELYS^ (Hattersley et al., 2016). Second, we found that several nucleoporins bearing potential membrane-binding domains were recruited downstream of MEL-28^ELYS^ to the interkinetic envelope. Our results suggested that these nucleoporins could recruit membranes necessary for interkinetic envelope assembly at the surface of chromosomes. The role of MEL-28^ELYS^ in nucleoporin recruitment to the interkinetic envelope could be direct or indirect. Indeed, during post-mitotic nuclear envelope reassembly, a key initial event is the dephosphorylation of nuclear envelope components, including lamins and nucleoporins, by protein phosphatases PP1 or PP2A (Hattersley et al., 2016; Mehsen et al., 2018; Steen et al., 2000). During meiosis I in *C. elegans* oocytes, MEL-28^ELYS^ is responsible for the docking of the catalytic subunit of PP1 on chromosomes (Hattersley et al., 2016). Thus, PP1 docked by MEL-28^ELYS^ on chromosomes could regulate the phosphorylation state of components essential for interkinetic envelope assembly, which would in turn promote their chromosomal recruitment.

The potential link we establish between nucleoporins bearing membrane-binding domains and interkinetic envelope assembly pertains to a non-conventional role of these nucleoporins outside their canonical function in the formation of nuclear pores. We indeed observed a complete lack of nuclear pores in the interkinetic envelope despite the presence of these nucleoporins. Consistent with this observation, we found that nucleoporins that normally belong to the same NPC sub- complex were not co-recruited to the interkinetic envelope. In a broader context, our findings imply that the conventional hierarchical relationships observed among nucleoporins within the same subcomplexes during post-mitotic nuclear pore assembly may not be preserved within the interkinetic envelope. We indeed found that NPP-21^TPR^ was absent despite the presence of NPP-7^NUP153^, which is both necessary and sufficient for its recruitment to NPCs during interphase (Hase and Cordes, 2003; Walther et al., 2001). While the recruitment of NPP-8^NUP155^ and NPP- 19^NUP35^ are interdependent at NPCs, we observed NPP-19^NUP35^ localized in the absence of NPP-8^NUP155^ at the interkinetic envelope (Rodenas et al., 2009). Moreover, nucleoporins essential for nuclear pore assembly (i.e., NPP-8^NUP155^) were even missing from the interkinetic envelope (Franz et al., 2005). The lack of NPCs in the interkinetic envelope is not surprising in light of their normal function in regulating transport between the physically segregated nucleoplasm and cytoplasm during interphase. The interkinetic envelope is a transient structure that only exists during the short transition period between meiosis I and II, and unlike the nuclear envelope, the interkinetic envelope never seals completely. During this stage, the condensed chromosomes are thus in contact with the cytoplasmic content, and can in theory freely exchange components.

Despite the lack of complete interkinetic envelope sealing, we nevertheless found that the structural integrity of the interkinetic envelope is functionally important. In the absence of the chromatin-binding protein BAF-1^BAF^, the interkinetic envelope was highly fenestrated and membrane fragments were observed between chromosomes of a given chromosomal set. This phenotype is reminiscent of the post-mitotic nuclear envelope defects, including envelope fragmentation and micronucleation, observed following BAF depletion in human tissue cultured cells and during *C. elegans* mitosis (Barger et al., 2023; Gorjánácz et al., 2007; Samwer et al., 2017). Recent studies have revealed that BAF not only plays a role in recruiting LEM-domain proteins for nuclear envelope assembly, as previously thought, but is also involved in DNA cross-bridging (Samwer et al., 2017). This function is carried out by BAF on chromosomes, where it creates a mechanically rigid surface of chromatin. This rigid chromatin surface restricts nuclear membranes to the chromosome surface and effectively prevents membrane fragmentation. Whether BAF-1^BAF^ functions by promoting chromosomal cohesion, targeting LEM-domain proteins, or through a combination of both mechanisms during interkinetic envelope assembly remains unclear. However, several lines of evidence suggest that both mechanisms may be involved. First, the presence of membrane fragments within chromosomal sets in the absence of BAF-1^BAF^—a phenotype never observed under normal conditions—implies that BAF-1^BAF^ may be crucial for excluding membranes from the spaces between chromosomes, suggesting its role in chromosomal cohesion. Second, our finding that depleting VRK-1^VRK1^, the kinase that negatively regulates BAF-1^BAF^ chromatin binding, produces the same fenestrated interkinetic envelope phenotype as BAF-1^BAF^ depletion but without interchromosomal membrane fragments, shows that these two phenotypes can be functionally separated. VRK-1^VRK1^ regulates BAF- 1^BAF^ chromatin binding, which recruits LEM domain proteins, such as LEM-2^LEMD2/3^ and EMR-1^Emerin^, to the chromosome surface. Therefore, VRK-1^VRK1^ and BAF-1^BAF^ depletion have opposite effects on LEM-domain protein chromosomal localization. Our observation that both the absence and over-recruitment of LEM-2^LEMD2/3^ is coupled to the same fenestrated interkinetic envelope phenotype could suggest that precise regulation of LEM domain protein localization is the essential factor for proper interkinetic envelope assembly. A key question is the potential function of the interkinetic envelope. While it may simply result from chromosomes remaining permanently condensed and exposed in the absence of spindle microtubules during interkinesis—allowing the transient recruitment of some inner nuclear envelope components and associated membranes—our findings suggest that the interkinetic envelope could play an active role in meiotic chromosome segregation. In the absence of BAF-1^BAF^, we observed fragmentation of the envelope, leading to accelerated and more extensive chromosome segregation compared to control oocytes. This suggests that the interkinetic envelope could act as a brake for meiotic chromosome segregation by mechanically constraining chromosomes and the pushing anaphase spindle. However, we believe that the observed phenotype may underestimate the true importance of the interkinetic envelope. Indeed, upon depletion of BAF-1^BAF^, although the interkinetic envelope becomes abnormally fenestrated, it still surrounds the surface of chromosomes, unlike the total absence of an envelope seen upon MEL-28^ELYS^ depletion. A striking observation is the abnormal spreading of chromosomes within each set in the absence of MEL-28^ELYS^, suggesting that the interkinetic envelope may play a role in keeping each chromosomal set tightly clustered to prevent the mixing of PB and MII chromosomes during segregation. Unfortunately, the severe chromosome segregation defects resulting from the lack of PP1-mediated kinetochore disassembly during meiotic anaphase after MEL-28^ELYS^ depletion prevented us from selectively analyzing the contribution of the interkinetic envelope to chromosome segregation (Gomez-Saldivar et al., 2016; Hattersley et al., 2016). In turn, the contribution of the interkinetic envelope defects in the chromosome segregation phenotype upon MEL-28^ELYS^ depletion is hard to estimate. Thus far, we have been unable to recapitulate the complete absence of an interkinetic envelope, nor the chromosome segregation defects observed in the absence of MEL-28^ELYS^, using other experimental perturbations. Understanding the true function of the interkinetic envelope in meiotic chromosome segregation will require further investigation.

The scarcity of research on the transient phase of meiosis has left the existence of an interkinetic envelope in oocytes of species other than *C. elegans* largely unknown. Investigating interkinesis in oocytes of other species will be an interesting avenue for future studies.

## MATERIALS AND METHODS

### Maintenance of *C. elegans* lines

The worm lines used in this study are listed in Supplementary Table 1. The worms were maintained on plates containing nematode growth medium (NGM) agar seeded with OP50 *E. coli* bacteria at 23 °C. All worms analyzed were hermaphrodites.

### RNA Interference

Double-stranded RNAs (dsRNAs) used in this study are listed in Supplementary Table 2. They were synthesized using the primers and templates indicated in the same table. PCR products were purified (PCR purification kit, Qiagen) and used as templates for T3 and T7 transcription reactions (Megascript, Invitrogen, #AM1334 for T7 and #AM1338 for T3). The produced RNAs were purified (MEGAclear kit, Invitrogen, #AM1908) and then hybridized by incubation at 68 °C for 10 minutes, followed by 37 °C for 30 minutes. L4 stage hermaphrodites were injected with dsRNAs at the specified concentrations and incubated at 20 °C for 44-48 hours before imaging.

### Auxin-Induced Degradation

The strain expressing endogenously tagged baf-1 (PHX2768) with an auxin-inducible degron (mAID) was first crossed with the strain CA1199, expressing a TIR1 transgene (sun-1p::TIR1::mRuby::sun-1 3’UTR + Cbr-unc-119(+)) (Zhang et al., 2015) under the control of the germline specific sun-1 promoter, and then with the GFP::TBA-2^α-tubulin^ and mCherry::H2B, under the control of the germline specific mex- 5 promoter, -expressing worms from the strain JDU233. Full BAF-1 depletion was achieved by combining RNAi-mediated depletion of the *baf-1* mRNA and auxin-induced degradation of the BAF-1 protein. Briefly, 32 hours after injection of dsRNA targeting *baf-1* in JDU647, worms were incubated for 16 hours on NGM agar plates seeded with OP50 *E. coli* bacteria containing 4 mM auxin.

### Oocyte Time-lapse imaging

Adult worms were dissected in 5 µL of meiosis medium (0.5 mg/mL Inulin, 25 mM HEPES, 60% Leibovitz L-15 medium, and 20% fetal bovine serum). Imaging was conducted at 23 °C using the CherryTemp temperature control system (CherryBiotech). All acquisitions were performed with a Nikon Ti-E inverted microscope, equipped with a CSU-X1 spinning disk confocal head (Yokogawa), an emission filter wheel, and a coolSNAP HQ2 CCD camera (Photometrics Scientific). Stage control and focus correction during acquisition were conducted using the PZ- 2000 XYZ piezo motor from Applied Scientific Instrumentation (ASI). Movies were acquired with 2 x 2 binning using a Nikon CFI APO S 60x/NA1.4 oil immersion objective. For all movies, 4 Z-stack planes, separated by 2 µm, were acquired every 20 seconds. Acquisition parameters were controlled using Metamorph 7 software (Molecular Devices, RRID:SCR_002368). For nocodazole treatment, adult worms were dissected in meiosis medium supplemented with 100 ng/µL nocodazole (Sigma, #M1404).

### Serial Block-Face Scanning Electron Microscopy (SBF-SEM)

After dissecting worms in 5 µL of meiosis medium, oocytes were transferred and packed into nitrocellulose capillary tubes with an inner diameter of 200 μm (Leica Microsystems, 16706869). The tube was sealed using the flat top edge of the scalpel. Oocytes enclosed in the capillary tubes were maintained on a glass slide in a droplet of meiosis medium and recorded by video-microscopy under the spinning disc confocal microscope as described above. Once in interkinesis, 15 µL of fixative medium (1% glutaraldehyde, 2% formaldehyde in 1x PBS) was added to the 5 µL of meiosis medium containing the capillary tube. Then, each capillary containing an oocyte was transferred into a 1.5 mL Eppendorf tube containing 1 mL of fixative medium. The oocytes were subsequently incubated for 1 hour at room temperature and kept at 4 °C until further preparation (Deerinck et al., 2010). After three washes in 1x PBS, the oocytes were treated with 1% osmium tetroxide (OsO_4_), 1,5% potassium ferrocyanide in 1X PBS at 4 °C for an hour. They were then incubated in a 1% thiocarbohydrazide (TCH) solution in water for 20 minutes at room temperature. Subsequently, they were treated with 2% aqueous OsO_4_ for 30 minutes at room temperature, before an overnight incubation at 4 °C in 1% uranyl acetate in water. The following day, the samples were subjected to Walton’s lead aspartate block staining (Walton, 1979) and placed in an oven at 60 °C for 30 minutes. The samples were then dehydrated in gradual ethanol concentrations (20%, 30%, 50%, 70%, 90%, and 100%) for 10 minutes each at room temperature on a wheel. The samples were infiltrated with a low-viscosity Agar resin (Agar Scientific Ltd) at 30% for 1 hour, then at 50% for 2 hours, at 75% for 2 hours, and finally at 100% overnight. The resin was then replaced, and the samples were re-included for 3 hours before being mounted and polymerized for 18 hours at 60 °C. The samples, permeated with 100% resin, were embedded in a flat layer of resin and then polymerized at 60 °C for 18 hours. The polymerized blocks were mounted on special aluminum pins for SBF-SEM imaging (FEI Microtome 8mm SEM Stub, Agar Scientific), with a two-part silver epoxy conduction kit (EMS, 190215). The samples mounted on aluminum pins were cut and inserted into a TeneoVS scanning electron microscope (Thermo Fisher Scientific). The acquisitions were carried out with a beam energy of 2 kV, 200 pA, in LowVac mode at 40 Pa, a pixel dwell time of 1 µs, and serial-sections of 30 nm and imaging was performed. The IMOD software (RRID:SCR_003297) was then used for stack reconstructions and segmentation (Kremer et al., 1996).

### Immunofluorescence

Ten to fifteen adult worms were dissected in 3.5 µL of meiosis medium on poly-l-lysine-coated slides (1 mg/mL in PBS, Sigma P-1524). The slides were covered with a 12 x 12 mm coverslip and snap-frozen in liquid nitrogen. The oocytes were then fixed in 100% methanol for 20 minutes at -20 °C. After two 10-minute washes in 1X PBS, the oocytes were blocked in an antibody diluent solution (AbDil containing 4% bovine serum albumin and 0.1% Triton in PBS) for an hour at room temperature in a humid chamber. The samples were subsequently incubated overnight at 4 °C in a primary antibody solution.(Supplementary table 3). After two washes in AbDil, the samples were incubated for an hour at room temperature with 1: 100 secondary antibodies. After two washes in AbDil, DNA was counterstained with 2 μg/mL Hoechst 33342 for 10 minutes. They were then washed twice with 1X PBS + 0.1% Triton X- 100 and once with 1X PBS. Samples were mounted between the glass slide and an 18 x 18 mm #1.5 coverslip in mounting medium (0.5% p-phenylenediamine in 90% glycerol and 20mM Tris pH 8.8) and stored at -20 °C. Acquisitions were carried out using the same microscope as above except without binning and a Nikon APO λS 100 x/1.45 oil objective. All immunofluorescence images are maximum projections of Z-stacks with Z-plans acquired every 0.2 μm.

### Embryonic viability assays and brood size

Embryonic viability assays were performed at 23 °C. For each condition, L4 stage worms were singled onto plates to lay embryos. Each day, for five consecutive days, the worms were transferred to new plates. Embryos were scored after transferring the parent worms and again 24 hours later to count the larvae. Embryonic viability was determined as the percentage of live embryos found within the progeny, and brood size was measured as the sum of the larvae.

### Image analyses

Image analyses were performed on maximum projections using the Fiji software (Schindelin et al., 2012, RRID:SCR_002285), and following the methods described in (Hattersley et al., 2018). Briefly, normalized intensities, in Fig. 1 E, Fig. 3 B, Fig. 4 C, D, Fig. 5 E, F, Fig. 7 A, B; Fig. 8, Fig. S1 D, Fig. S2 D, G, Fig. S3 E, Fig. S4, Fig. S5 A were quantified by drawing a rectangular box around the MII chromosomal set and measuring its area (A_a_) and integrated intensity (I_a_) at each time point. The background intensity was quantified by measuring the area (A_b_) and the integrated intensity of an expanded rectangle (5 pixels on every side) (I_b_) around the MII chromosomal set. The background signal (B_s_) corresponds to the difference of the signal and area between the expanded rectangle and the original one B_s_ = (I_b_ - I_a_)/((A_b_ - A_a_)/ A_a_). Finally, the normalized integrated intensity over the background corresponds to the difference between the background signal and the intensity of the original signal: (I_a_ – B_s_)/A_a_. Chromosome segregation in Fig. 3 D and S5 A was quantified by measuring the distance between the inner surfaces of the chromosome sets over time.

### Graphs and Statistics

GraphPad Prism 8 (RRID:SCR_002798) was used to generate all graphs and perform statistical tests as indicated in the figure legends.

## SUMMARY OF SUPPLEMENTAL MATERIAL

5 supplementary figures

10 supplementary videos

3 supplementary tables

## DATA AVAILABILITY

All data supporting the findings of this study are available within the paper and its Supplementary Information.

## ACKNOWLEDGEMENTS

We thank all members of the Dumont lab for support and advice. We are grateful to Patricia Moussounda, Clarisse Picard, and Téo Bitaille for providing technical support. We thank Anjon Audhya for the generous gift of antibodies.

We acknowledge the ImagoSeine core facility of Institut Jacques Monod, member of France-BioImaging (ANR-10-INBS-04) and IBiSA, with the support of Labex "Who Am I", Inserm Plan Cancer, Region Ile-de-France and Fondation Bettencourt Schueller. This work was supported by CNRS and University Paris Cité, by NIH R01GM117407 and R01GM130764 (J.C. Canman), by grant from the Spanish Agencia Estatal de Investigación and the European Regional Development Fund PID2019-105069GB-I00 (P. Askjaer), by 4th-year Ph.D. fellowship from the Ligue Nationale Contre le Cancer RS30J21DOC17_ELMOSSADEQ (L. El Mossadeq), and by grant from the European Research Council ERC-CoG ChromoSOMe 819179 (J. Dumont).

## AUTHOR CONTRIBUTIONS

Conceptualization: LEM, JD

Methodology: LEM, LB, RL, JMV, LP, PA, JCC, JD

Investigation: LEM, LB, JD Visualization: LEM, RL, JD Funding acquisition: JD Project administration: JD Supervision: JMV, JD

Writing – original draft: LEM, JD

Writing – review & editing: LEM, JCC, JD

## VIDEO LEGENDS

**Video 1: Ultrastructure of the interkinetic envelope throughout anaphase I and interkinesis.** 3-dimensional reconstructions centered on chromosomes of mid-anaphase I (left), mid-interkinesis (center) and late interkinesis (right) oocytes.

Chromosomes in magenta, membranes in contact with chromosomes in green, and plasma membrane in gray. Scale bar, 1 µm.

**Video 2: LEM-2^LEMD2/3^ localization in meiosis I and II.** Time-lapse imaging of an oocyte expressing mCherry::H2B (magenta) and GFP::LEM-2^LEMD2/3^ (green) during the meiotic and first mitotic division. Timings indicated are from anaphase I onset. Scale bar, 5 µm.

**Video 3: The interkinetic envelope contains inner, but lacks outer, nuclear membrane proteins.** Time-lapse imaging of oocytes expressing mCherry::H2B (magenta) and either GFP::EMR-1^Emerin^, GFP::BAF-1^BAF^, GFP::LMN-1^Lamin^ ^A^, SUN- 1^SUN1^::GFP, GFP::ZYG-12, GFP::SP12 or GFP::RAMP4 during meiosis I and II.

Timings indicated are from anaphase I onset. Scale bar, 5 µm.

**Videos 4: The interkinetic envelope is not connected to the endoplasmic reticulum.** 3-dimensional reconstructions centered on chromosomes of mid-anaphase I (left) and late interkinesis (right) oocytes. Chromosomes in magenta, membranes in contact with chromosomes in green, plasma membrane in gray, eggshell in gold, and endoplasmic reticulum in blue. Scale bar, 1 µm.

**Video 5: BAF-1^BAF^ and VRK-1^vrk1^ are essential for interkinetic envelope integrity.** 3-dimensional reconstructions centered on chromosomes of control (AID::BAF-1^BAF^, No auxin, No RNAi) (top), BAF-1^BAF^-depleted (AID::BAF-1^BAF^, 4 mM auxin, *baf- 1(RNAi)*) (middle), and VRK-1^vrk1^-depleted (*vrk-1(RNAi)*) (bottom) oocytes.

Chromosomes in magenta, membranes in contact with chromosomes in green, and plasma membrane in gray. Scale bar, 1 µm.

**Video 6: BAF-1^BAF^ is essential for LEM-2^LEMD2/3^ localization in interkinesis.** Time-lapse imaging of oocytes expressing mCherry::H2B (magenta) and GFP::LEM- 2^LEMD2/3^ (green) during meiosis I and II in the indicated conditions. Timings indicated are from anaphase I onset. Scale bar, 5 µm.

**Video 7: MEL-28^ELYS^ is essential for LEM-2^LEMD2/3^ localization in interkinesis.** Time-lapse imaging of oocytes expressing mCherry::H2B (magenta) and GFP::LEM- 2^LEMD2/3^ (green) during meiosis I and II in the indicated conditions. Timings indicated are from anaphase I onset. Scale bar, 5 µm.

**Video 8: MEL-28^ELYS^ is required for interkinetic envelope integrity.** 3-dimensional reconstructions centered on chromosomes of a control (top) and a MEL-28^ELYS^- depleted (bottom) oocytes. Chromosomes in magenta, membranes in contact with chromosomes in green, membranes distant from chromosomes in orange, vesicles in yellow, mitochondria in purple, endoplasmic reticulum in blue, and plasma membrane in gray. Scale bar, 1 µm.

**Video 9: Nucleoporins with a membrane-binding domain could contribute to interkinetic envelope integrity.** Time-lapse imaging of oocytes expressing mCherry::H2B (magenta) and GFP::LEM-2^LEMD2/3^ (green) during meiosis I and II in the indicated conditions. Timings indicated are from anaphase I onset. Scale bar, 5 µm.

**Video 10: The localization of nucleoporins with membrane-binding domains is partially or entirely dependent on MEL-28^ELYS^.** Time-lapse imaging of oocytes expressing mCherry::H2B (magenta) and either GFP-tagged NPP-6^NUP160^, NPP- 15^NUP133^, NPP-25^TMEM33^, or NPP-7^NUP153^ (green) during meiosis I and II in the indicated conditions. Timings indicated are from anaphase I onset. Scale bar, 5 µm.

