## Supplementary Figures for "An interkinetic envelope surrounds chromosomes between meiosis I and II in C. elegans oocytes"

### Figure S1

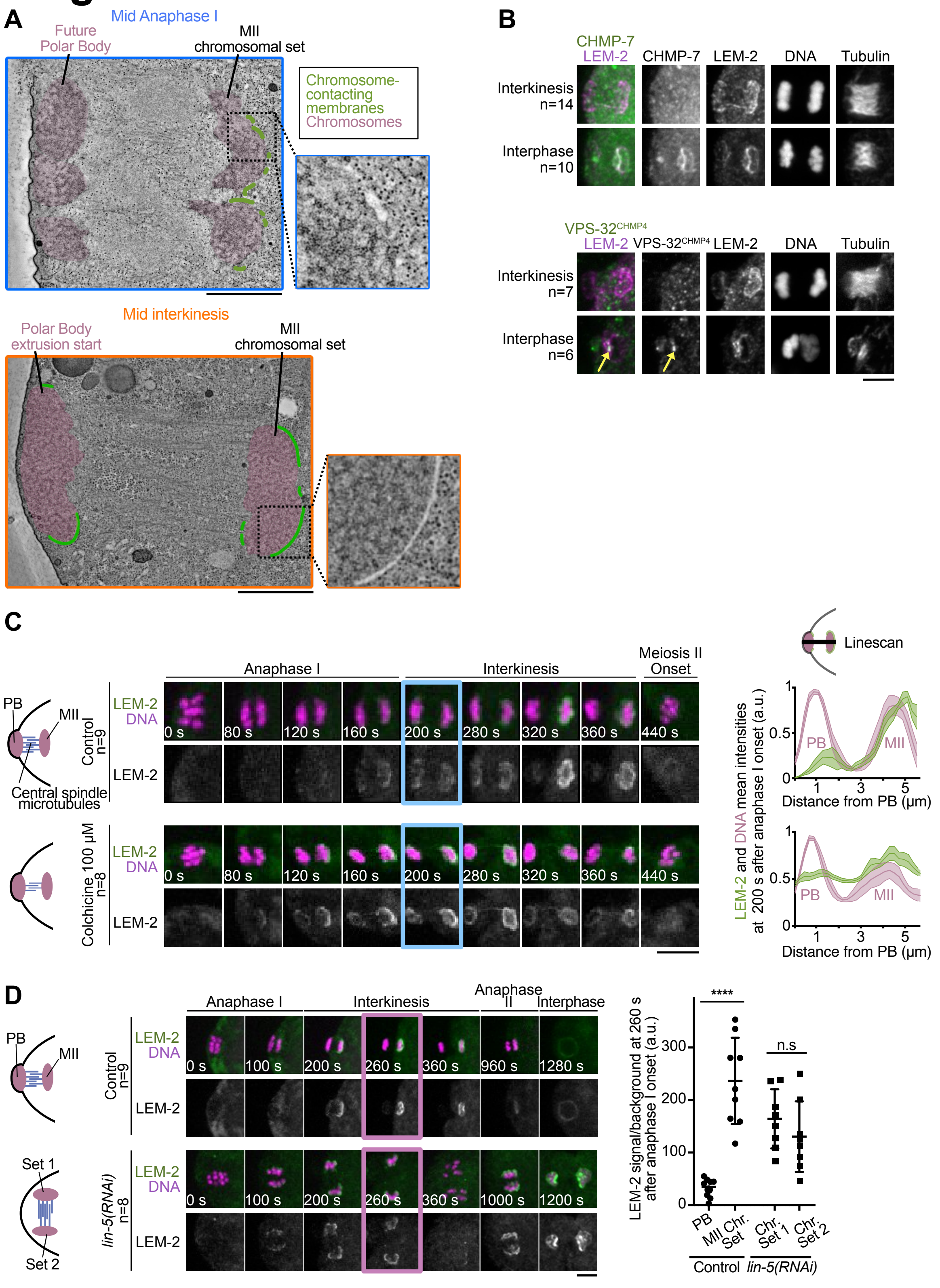
A
Mid Anaphase I
Future Polar Body
MII chromosomal set
Chromosome-contacting membranes
Chromosomes
Mid interkinesis
Polar Body extrusion start
MII chromosomal set
B
CHMP-7
LEM-2
CHMP-7
LEM-2
DNA
Tubulin
Interkinesis n=14
Interphase n=10
VPS-32CHMP4
LEM-2
VPS-32CHMP4
LEM-2
DNA
Tubulin
Interkinesis n=7
Interphase n=6
C
Linescan
Anaphase I
Interkinesis
Meiosis II Onset
PB
MII
Central spindle microtubules
Control n=9
LEM-2
DNA
LEM-2
0 s
80 s
120 s
160 s
200 s
280 s
320 s
360 s
440 s
Colchicine 100 µM n=8
LEM-2 and DNA mean intensities at 200 s after anaphase I onset (a.u.)
Distance from PB (µm)
D
Anaphase I
Interkinesis
Anaphase II
Interphase
Control n=9
lin-5(RNAi) n=8
Set 1
Set 2
0 s
100 s
200 s
260 s
360 s
960 s
1280 s
1000 s
1200 s
LEM-2 signal/background at 260 s after anaphase I onset (a.u.)
****
n.s
PB
MII Chr. Set
Chr. Set 1
Chr. Set 2
Control
lin-5(RNAi)

**Figure S1: The interkinetic envelope is never sealed. (A)** Electron tomography sections centered on chromosomes of mid-anaphase I (left) and mid interkinesis (right) oocytes showing each chromosome set in magenta and membranes in contact with chromosomes in green. Each section is accompanied (right) by a magnification of an ROI centered on a portion of the MII chromosomal set. Scale bar, 1 µm. **(B)** Representative images centered on chromosomes of fixed oocytes showing the immunolocalization of LEM-2$^{LEMD2/3}$, DNA, Tubulin and either CHMP-7$^{CHMP7}$ (top) or VPS-32$^{CHMP4}$ (bottom) in interphase and interkinesis. Yellow arrows indicate the focus of VPS-32$^{CHMP4}$ at the site of envelope sealing in interphase. Scale bar, 5 µm. **(C)** Left: Schematics of chromosomes and anaphase I central spindle microtubules in the indicated conditions. Middle: Representative time-lapse images centered on chromosomes of oocytes expressing mCherry::H2B (magenta) and GFP::LEM-2$^{LEMD2/3}$ (green) during meiosis I and II in the indicated conditions. Timings indicated at the bottom left corners of images are from anaphase I onset. The specific meiotic stage used for the quantification is highlighted in cyan. Scale bar, 5 µm. Right: Quantifications of LEM-2$^{LEMD2/3}$ and DNA mean intensities at 200 s after anaphase I onset in the indicated conditions. **(D)** Left: Schematics of chromosomes and spindle orientation in indicated conditions. Middle: Representative time-lapse images centered on chromosomes of oocytes expressing mCherry::H2B (magenta) and GFP::LEM-2$^{LEMD2/3}$ (green) during meiosis I and II in the indicated conditions. Timings indicated at the bottom left corners of images are from anaphase I onset. The timing used for the quantification is highlighted in pink. Scale bar, 5 µm. Right: Quantification of the normalized GFP::LEM-2$^{LEMD2/3}$ integrated intensity at 260 s after anaphase I onset. Error bars correspond to the standard deviation. Mann-Whitney test (**** $p < 0.0001$).

### Figure S2

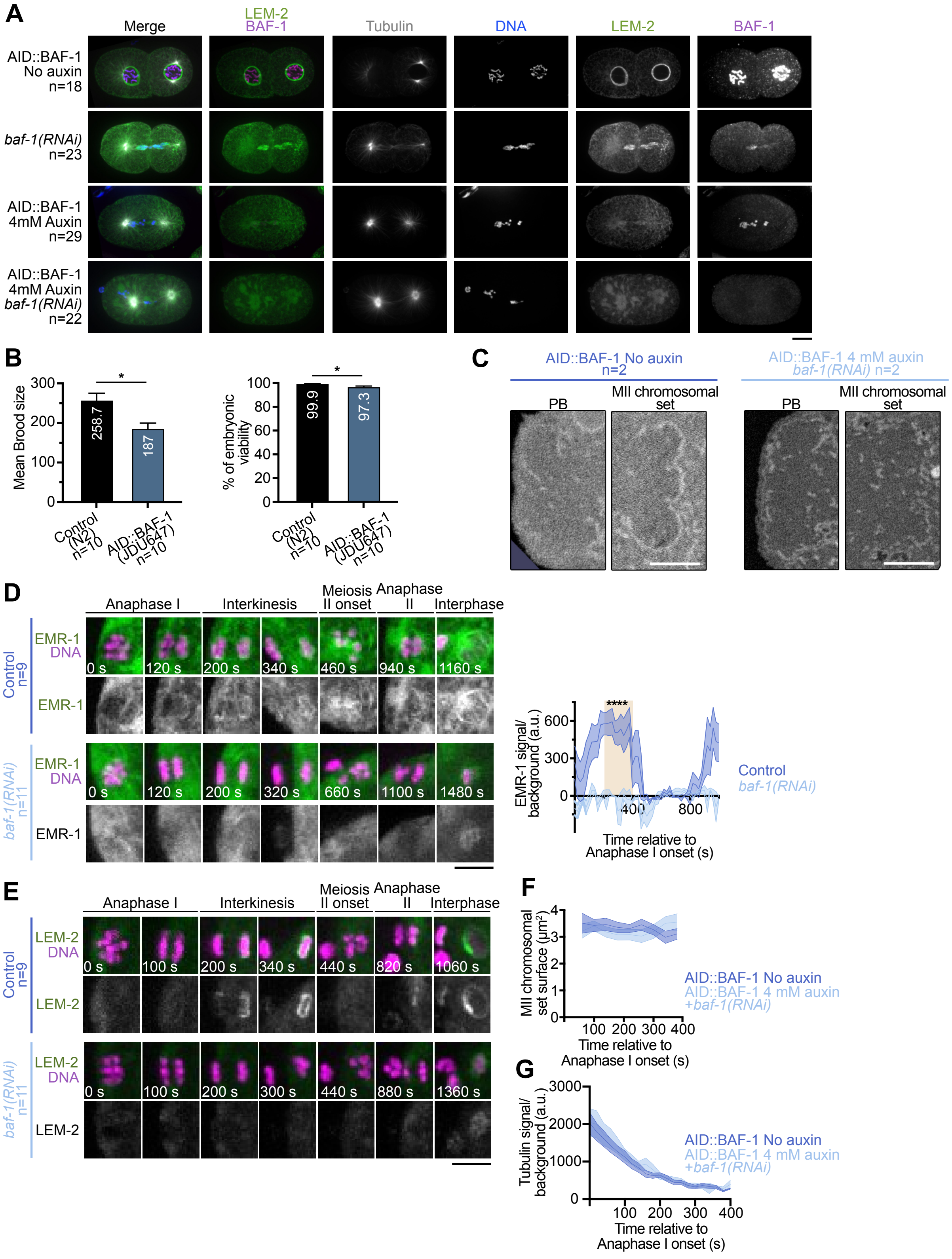
A
Merge
LEM-2
BAF-1
Tubulin
DNA
LEM-2
BAF-1
AID::BAF-1
No auxin
n=18
baf-1(RNAi)
n=23
AID::BAF-1
4mM Auxin
n=29
AID::BAF-1
4mM Auxin
baf-1(RNAi)
n=22
B
Mean Brood size
300
200
100
0
258.7
187
*
Control (N2) n=10
AID::BAF-1 (JDU647) n=10
% of embryonic viability
100
80
60
40
20
0
99.9
97.3
*
Control (N2) n=10
AID::BAF-1 (JDU647) n=10
C
AID::BAF-1 No auxin n=2
AID::BAF-1 4 mM auxin baf-1(RNAi) n=2
PB
MII chromosomal set
PB
MII chromosomal set
D
Anaphase I
Interkinesis
Meiosis II onset
Anaphase II
Interphase
Control n=9
EMR-1
DNA
0 s
120 s
200 s
340 s
460 s
940 s
1160 s
EMR-1
baf-1(RNAi) n=11
EMR-1
DNA
0 s
120 s
200 s
320 s
660 s
1100 s
1480 s
EMR-1
EMR-1 signal/ background (a.u.)
600
300
0
400
800
****
Time relative to Anaphase I onset (s)
Control
baf-1(RNAi)
E
Anaphase I
Interkinesis
Meiosis II onset
Anaphase II
Interphase
Control n=9
LEM-2
DNA
0 s
100 s
200 s
340 s
440 s
820 s
1060 s
LEM-2
baf-1(RNAi) n=11
LEM-2
DNA
0 s
100 s
200 s
300 s
440 s
880 s
1360 s
LEM-2
F
MII chromosomal set surface (µm²)
4
3
2
1
0
100
200
300
400
Time relative to Anaphase I onset (s)
AID::BAF-1 No auxin
AID::BAF-1 4 mM auxin
+baf-1(RNAi)
G
Tubulin signal/ background (a.u.)
3000
2000
1000
0
100
200
300
400
Time relative to Anaphase I onset (s)
AID::BAF-1 No auxin
AID::BAF-1 4 mM auxin
+baf-1(RNAi)

**Figure S2: BAF-1$^{BAF}$ is required for the recruitment of LEM-2$^{LEMD2/3}$ and EMR-1$^{Emerin}$ at the chromosome surface in interkinesis. (A)** Representative images centered on chromosomes of fixed embryos showing the immunolocalization of LEM-2$^{LEMD2/3}$, DNA, Tubulin and BAF-1$^{BAF}$ during pronuclei migration in control conditions, *baf-1(RNAi)* and/or 4 mM auxin treatment. Scale bar, 5 µm. **(B)** Quantifications of the mean brood size (left) and the percentage of embryonic viability (right) in N2 (control) and JDU647 (AID::BAF-1$^{BAF}$) strains. **(C)** 2-dimensional single sections of two ROIs centered on each chromosomal set of a control oocyte (AID::BAF-1$^{BAF}$, No auxin) (top) and a BAF-1$^{BAF}$-depleted oocyte (AID::BAF-1$^{BAF}$, 4 mM auxin, *baf-1(RNAi)*). Scale bar, 1 µm. **(D)** Left: Representative time-lapse images centered on chromosomes of oocytes expressing mCherry::H2B (magenta) and GFP::EMR-1$^{Emerin}$ (green) during meiosis I and II in the indicated conditions. Timings indicated at the bottom left corners of images are from anaphase I onset. Scale bar, 5 µm. Right: Quantification of the GFP::EMR-1$^{Emerin}$ integrated intensity normalized over background from anaphase I onset to interphase for the MII chromosomal set. Control in dark blue, *baf-1(RNAi)* in light blue. Error bars correspond to the standard error of the mean. The orange box indicates interkinesis. Mann-Whitney test on the mean value of GFP::EMR-1$^{Emerin}$ intensity in interkinesis (**** $p<0.0001$). **(E)** Representative time-lapse images centered on chromosomes of oocytes expressing mCherry::H2B (magenta) and GFP::LEM-2$^{LEMD2/3}$ (green) during meiosis I and II in the indicated conditions. Timings indicated at the bottom left corners of images are from anaphase I onset. Scale bar, 5 µm. **(F, G)** Quantifications of the surface of the MII chromosomal set (F) and of tubulin intensity (G) over time from anaphase I onset in control oocytes (dark blue) and BAF-1$^{BAF}$-depleted oocytes (light blue). Error bars correspond to the standard error of the mean.

### Figure S3

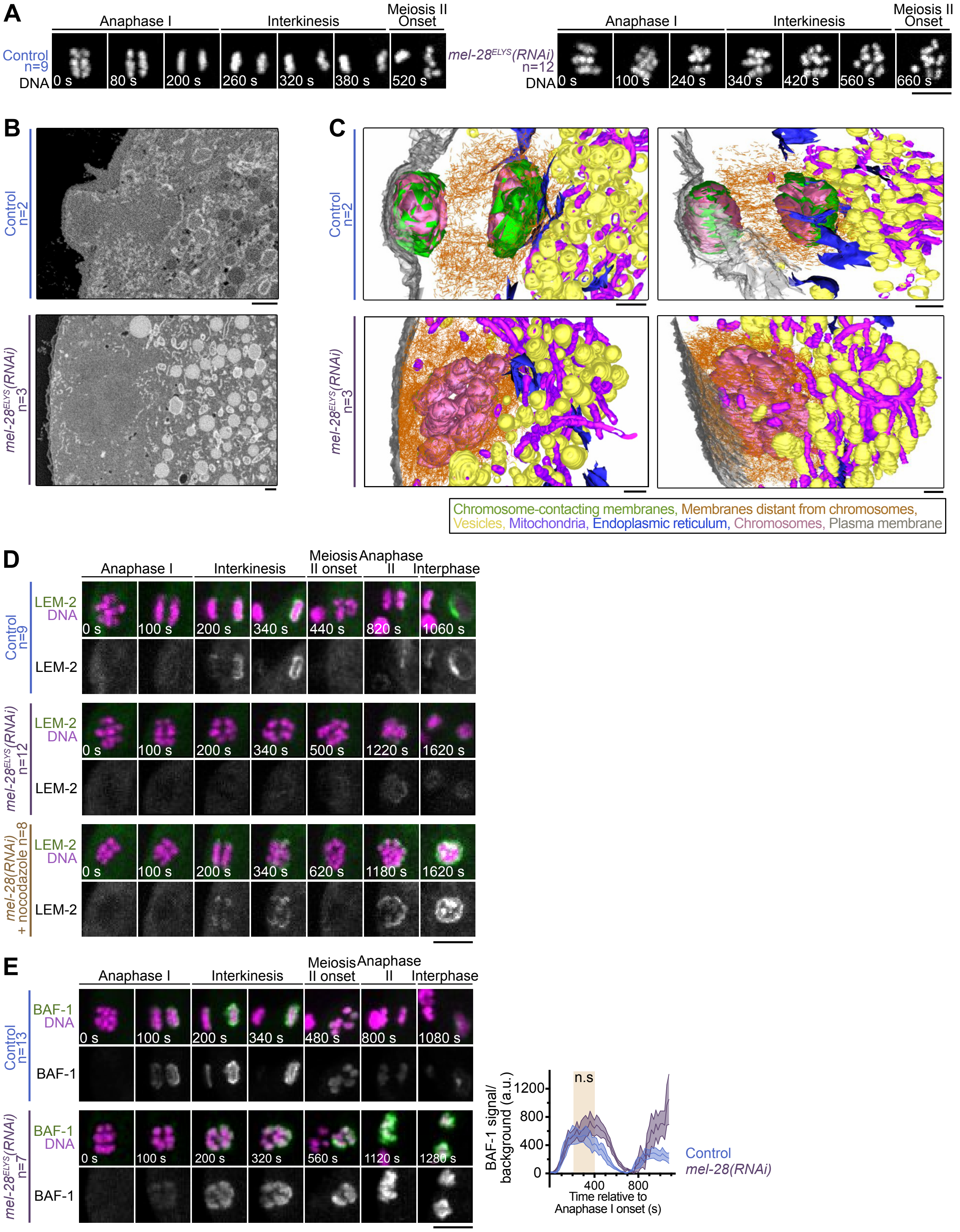
A
Anaphase I
Interkinesis
Meiosis II Onset
Control n=9
DNA
0 s
80 s
200 s
260 s
320 s
380 s
520 s
mel-28ELYS(RNAi) n=12
0 s
100 s
240 s
340 s
420 s
560 s
660 s
B
Control n=2
mel-28ELYS(RNAi) n=3
C
Control n=2
mel-28ELYS(RNAi) n=3
Chromosome-contacting membranes, Membranes distant from chromosomes, Vesicles, Mitochondria, Endoplasmic reticulum, Chromosomes, Plasma membrane
D
Anaphase I
Interkinesis
Meiosis II onset
Anaphase II
Interphase
Control n=9
LEM-2 DNA
0 s
100 s
200 s
340 s
440 s
820 s
1060 s
LEM-2
mel-28ELYS(RNAi) n=12
LEM-2 DNA
0 s
100 s
200 s
340 s
500 s
1220 s
1620 s
LEM-2
mel-28(RNAi) + nocodazole n=8
LEM-2 DNA
0 s
100 s
200 s
340 s
620 s
1180 s
1620 s
LEM-2
E
Anaphase I
Interkinesis
Meiosis II onset
Anaphase II
Interphase
Control n=13
BAF-1 DNA
0 s
100 s
200 s
340 s
480 s
800 s
1080 s
BAF-1
mel-28ELYS(RNAi) n=7
BAF-1 DNA
0 s
100 s
200 s
320 s
560 s
1120 s
1280 s
BAF-1
n.s
BAF-1 signal/ background (a.u.)
1200
800
400
0
400
800
Time relative to Anaphase I onset (s)
Control
mel-28(RNAi)

**Figure S3: MEL-28$^{ELYS}$ is required for the interkinetic envelope integrity. (A)** Representative time-lapse images centered on chromosomes of oocytes expressing mCherry::H2B (gray) during anaphase I and interkinesis in the indicated conditions. Timings indicated at the bottom left corners of images are from anaphase I onset. Scale bar, 5 µm. **(B)** 2-dimensional single sections of two ROIs centered on each chromosomal set of a control (top) and a MEL-28$^{ELYS}$-depleted (bottom) oocyte. Scale bars, 1 µm. **(C)** 3-dimensional reconstructions centered on chromosomes of a control oocyte (top) and a MEL-28$^{ELYS}$-depleted oocyte (bottom) viewed from two different angles. Scale bar, 1 µm. Chromosomes in magenta, membranes in contact with chromosomes in green, membranes distant from chromosomes in orange, vesicles in yellow, mitochondria in purple, endoplasmic reticulum in blue, and plasma membrane in gray. Scale bars, 1 µm. **(D)** Representative time-lapse images centered on chromosomes of oocytes expressing mCherry::H2B (magenta) and GFP::LEM-2$^{LEMD2/3}$ (green) during meiosis I and II in the indicated conditions. Timings indicated at the bottom left corners of images are from anaphase I onset. Scale bar, 5 µm. **(E)** Left: Representative time-lapse images centered on chromosomes of oocytes expressing mCherry::H2B (magenta) and GFP::BAF-1$^{BAF}$ (green) during meiosis I and II in the indicated conditions. Timings indicated at the bottom left corners of images are from anaphase I onset. Scale bar, 5 µm. Right: Quantification of the GFP::BAF-1$^{BAF}$ integrated intensity normalized over background from anaphase I onset to interphase for the MII chromosomal set. Control in dark blue, *mel-28(RNAi)* in purple. Error bars correspond to the standard error of the mean. The orange box indicates interkinesis. Mann-Whitney test on the mean value of GFP::BAF-1$^{BAF}$ intensity in interkinesis.

### Figure S4

A

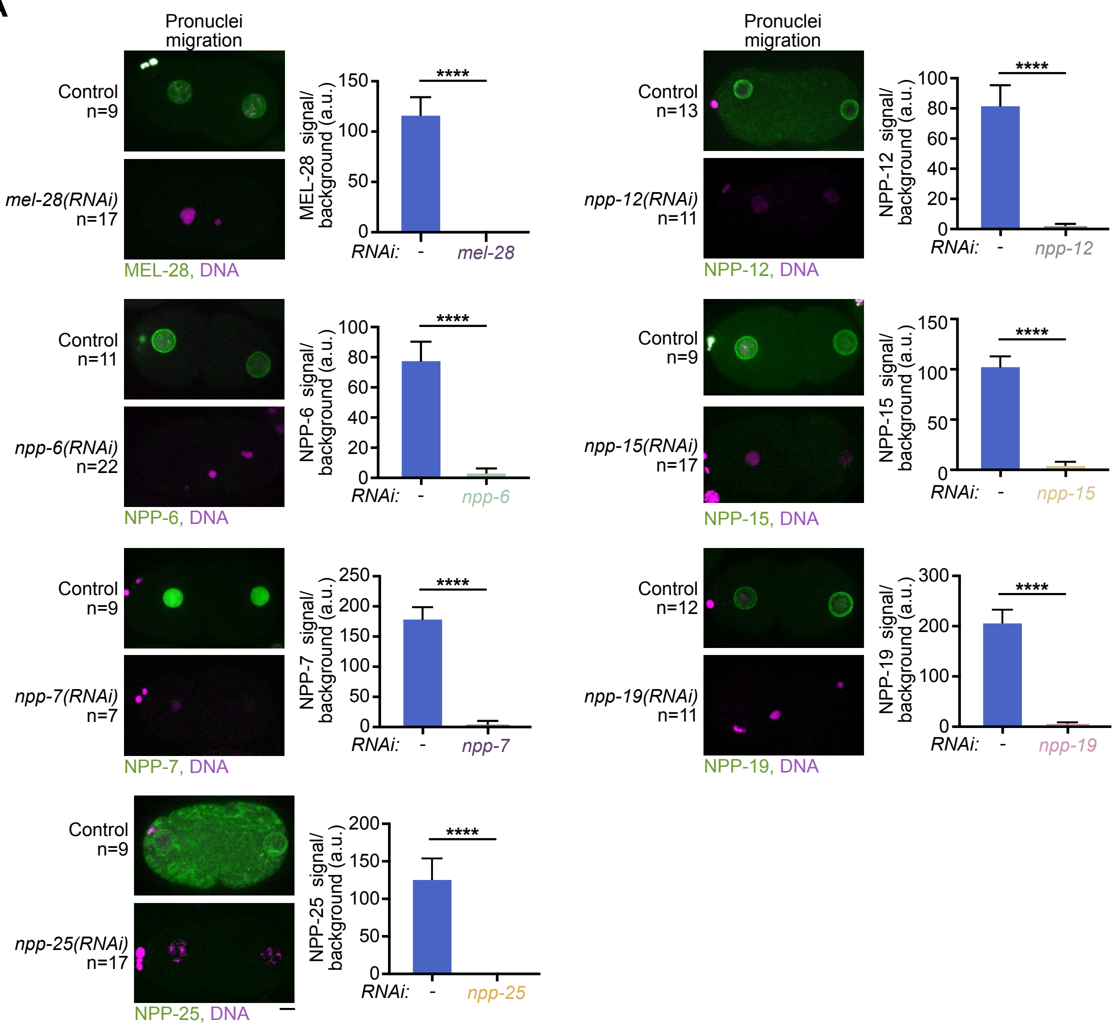

Pronuclei
migration
Control
n=9
mel-28(RNAi)
n=17
MEL-28, DNA
MEL-28 signal/
background (a.u.)
150
100
50
0
****
RNAi:
-
mel-28
Control
n=11
npp-6(RNAi)
n=22
NPP-6, DNA
NPP-6 signal/
background (a.u.)
100
80
60
40
20
0
npp-6
Control
n=9
npp-7(RNAi)
n=7
NPP-7, DNA
NPP-7 signal/
background (a.u.)
250
200
150
100
50
0
npp-7
Control
n=9
npp-25(RNAi)
n=17
NPP-25, DNA
NPP-25 signal/
background (a.u.)
200
150
100
50
0
npp-25
Pronuclei
migration
Control
n=13
npp-12(RNAi)
n=11
NPP-12, DNA
NPP-12 signal/
background (a.u.)
100
80
60
40
20
0
npp-12
Control
n=9
npp-15(RNAi)
n=17
NPP-15, DNA
NPP-15 signal/
background (a.u.)
150
100
50
0
npp-15
Control
n=12
npp-19(RNAi)
n=11
NPP-19, DNA
NPP-19 signal/
background (a.u.)
300
200
100
0
npp-19

**Figure S4 : Depletion efficiency of various nucleoporins by RNAi. (A)** Left: Representative images centered on chromosomes of embryos expressing mCherry::H2B (magenta) and either GFP-tagged MEL-28$^{ELYS}$, NPP-6$^{NUP160}$, NPP-15$^{NUP133}$, NPP-25$^{TMEM33}$, NPP-12$^{NUP210}$, NPP-19$^{NUP35}$ or NPP-7$^{NUP153}$ (green) during pronuclei migration in the indicated conditions. Scale bar, 5 µm. Right: Quantification of the normalized GFP-tagged MEL-28$^{ELYS}$, NPP-6$^{NUP160}$, NPP-15$^{NUP133}$, NPP-25$^{TMEM33}$, NPP-12$^{NUP210}$, NPP-19$^{NUP35}$ or NPP-7$^{NUP153}$ integrated intensity in an ROI centered around the maternal and paternal pronuclei in the indicated conditions. Error bars correspond to the standard error of the mean. Mann-Whitney test (**** $p<0.0001$).

### Figure S5

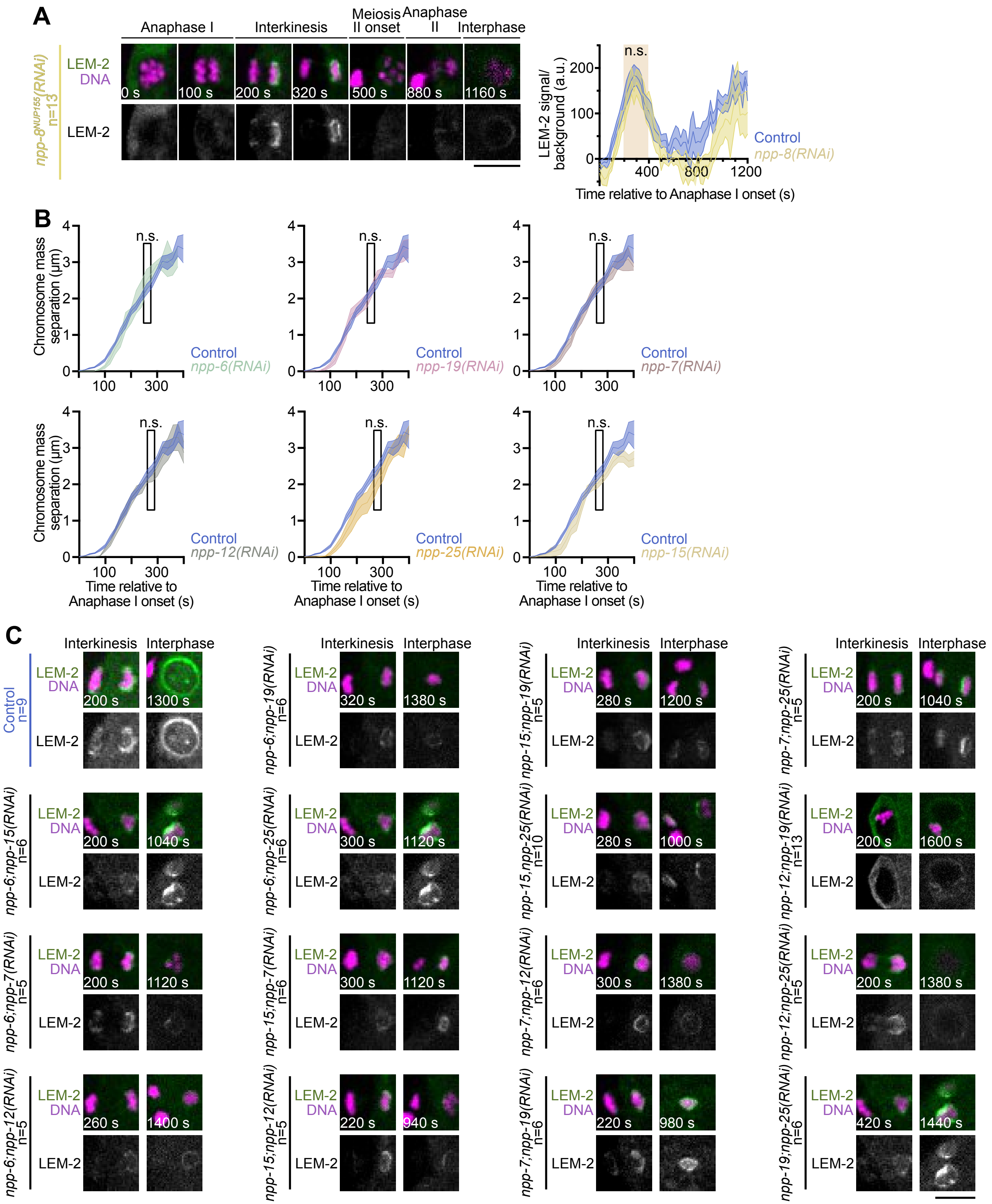
A
Anaphase I
Interkinesis
Meiosis II onset
Anaphase II
Interphase
npp-8NUP155(RNAi) n=13
LEM-2
DNA
0 s
100 s
200 s
320 s
500 s
880 s
1160 s
LEM-2 signal/ background (a.u.)
n.s.
200
100
0
400
800
1200
Control
npp-8(RNAi)
Time relative to Anaphase I onset (s)
B
Chromosome mass separation (µm)
n.s.
Control
npp-6(RNAi)
npp-19(RNAi)
npp-7(RNAi)
npp-12(RNAi)
npp-25(RNAi)
npp-15(RNAi)
Time relative to Anaphase I onset (s)
C
Interkinesis
Interphase
Control n=9
npp-6;npp-19(RNAi) n=6
npp-15;npp-19(RNAi) n=5
npp-7;npp-25(RNAi) n=5
npp-6;npp-15(RNAi) n=6
npp-6;npp-25(RNAi) n=6
npp-15,npp-25(RNAi) n=10
npp-12;npp-19(RNAi) n=13
npp-6;npp-7(RNAi) n=5
npp-15;npp-7(RNAi) n=6
npp-7;npp-12(RNAi) n=6
npp-12;npp-25(RNAi) n=5
npp-6;npp-12(RNAi) n=5
npp-15;npp-12(RNAi) n=5
npp-7;npp-19(RNAi) n=6
npp-19;npp-25(RNAi) n=6

**Figure S5: (A)** Left: Representative time-lapse images centered on chromosomes of oocytes expressing mCherry::H2B (magenta) and GFP::LEM-2$^{LEMD2/3}$ (green) during meiosis I and II in the indicated conditions. Timings indicated at the bottom left corners of images are from anaphase I onset. Scale bar, 5 µm. Right: Quantification of the GFP::LEM-2$^{LEMD2/3}$ integrated intensity normalized over background from anaphase I onset to interphase for the MII chromosomal set. Control in dark blue, *npp-8(RNAi)* in yellow. Error bars correspond to the standard error of the mean. The orange box indicates interkinesis. Mann-Whitney test on the mean value of GFP::LEM-2$^{LEMD2/3}$ intensity in interkinesis. **(B)** Quantification of the distance between the two sets of segregating chromosomes over time from anaphase I onset. Control in dark blue, *npp-6$^{NUP160}$(RNAi)* in turquoise, *npp-25$^{TMEM33}$(RNAi)* in dark orange, *npp-15$^{NUP133}$(RNAi)* in light orange, *npp-12$^{NUP210}$(RNAi)* in gray, *npp-19$^{NUP53}$(RNAi)* in pink and *npp-7$^{NUP153}$(RNAi)* in light brown. Mann-Whitney test on the mean distance between the segregating chromosomal sets during interkinesis in both conditions (** $p < 0.01$). **(C)** Representative time-lapse images centered on chromosomes of oocytes expressing mCherry::H2B (magenta) and GFP::LEM-2$^{LEMD2/3}$ (green) during interkinesis and interphase in the indicated conditions. Timings indicated at the bottom left corners of images are from anaphase I onset. Scale bar, 5 µm.
