## Supplementary Table 1 for "An interkinetic envelope surrounds chromosomes between meiosis I and II in C. elegans oocytes"

| Strain | Genotype | Source |
| --- | --- | --- |
| BN47 | npp-19(tm2886) II; ltIs37[Ppie-1::mCherry::his-58; unc-119 (+)] IV; bqIs07[unc-119(+); Ppie-1::LAP::npp-19] ? | Ródenas et al. 2009 |
| BN68 | bqIs51[Ppie-1::gfp::npp-5]; ltIs37[Ppie-1::mCherry::his-58] | Ródenas et al. 2012 |
| BN245 | ltIs37[Ppie-1::mCherry::his-58;unc-119(+)]IV;ojIs1[unc-119(+)Ppie 1::GFP::tbb-2]V(?) and/or ltIs24[pAZ132;pie-1/GFP::tba-2+unc-119(+)]? | Morales-Martínez et al., 2015 |
| BN246 | lem-2(tm1582)II; ltIs37[Ppie-1::mCherry::his-58; unc-119(+)] IV; ojIs1[unc-119(+) Ppie-1::GFP::tbb-2] V (?) and/or ltIs24[pAZ132; pie-1/GFP::tba-2 + unc-119(+)] ? | Morales-Martínez et al., 2015 |
| BN591 | vrk-1(ok1181) II; unc-119(ed3) his-72(cp10[his-72::gfp+ LoxP unc-119(+) LoxP]) III; bqSi171[p961(unc-119(+) Pvrk-1::vrk-1::mCherry)] IV | Dobrzynska et al., 2016 |
| BN593 | dpy-10(cn64) npp-21(bq1[npp-21::gfp]) II; ltIs37[Ppie-1::mCherry::his-58; unc-119 (+)] IV | This study |
| BN740 | npp-24(bq15[G>F>P::npp-24]) bqSi189[pBN13(unc-119(+) Plmn-1::mCherry::his-58)] II; may carry unc-119(ed3) or unc-119(ed9) III | Thomas, Askjaer, et Seydoux 2022 |
| BN1082 | npp-2(bq38[G>F>P::npp-2]) I; bqSi189[pBN13(unc-119(+) lmn-1p::mCherry::his-58)] II | This study |
| BN1112 | bqSi189[pBN13(unc-119(+) lmn-1p::mCherry::his-58)] II; npp-25(bq41[npp-25::G>F>P]) III | This study |
| JDU233 | ijmSi63 [pJD520; mosII_5'mex-5_GFP::tba-2; mCherry::his-11; cb-unc-119(+)] II; unc-119(ed3) III? | Pitayu-Nugroho, L., Aubry et al. 2023 |
| JDU424 | ijmSi49 [pJD479_pJD348_Mos1_Pmex-5_GFP::lmn-1_3'UTRtbb-2]I; ijmSi31 [pJD446_pJD362_Mos2_Pmex-5_mCherry::his11_3'UTRtbb-2]II; unc-119(ed3) III? | This study |
| JDU455 | ijmSi47 [pJD478_pJD348_Mos1_Pmex-5_GFP::emr-1_3'UTRtbb-2]I; ijmSi31 [pJD446_pJD362_Mos2_Pmex-5_mCherry::his11_3'UTRtbb-2]II; unc-119(ed3) III? | This study |
| JDU456 | [WRM0636A_D05(pRedFlp-Hgr)::unc-119-Nat(npp-11(37836)::2XTY1-eGFP-3XFlag)dFRT]?; ijmSi31 [pJD446_pJD362_Mos2_Pmex-5_mCherry::his11_3'UTRtbb-2]II; unc-119(ed3) III? | This study |
| JDU461 | ijmSi31 [pJD446_pJD362_Mos2_Pmex-5_mCherry::his11_3'UTRtbb-2]II; baf-1(bq12[GFP::baf-1]), unc-119(ed3) III? | This study |
| JDU489 | ttTi5605 jfSi1[Psun-1::GFP cb-unc-119(+)] II; unc-119(ed3) III?; ltIs37 [pAA64; pie-1/mCHERRY::his-58; unc-119 (+)] IV; sun-1(ok1282)V/nT1[qIs51](IV;V). | This study |
| JDU509 | ojIs9[zyg-12::GFP + unc-119(+)]?; unc-119(ed3) III?; ltIs37[pAA64; pie-1/mCHERRY::his-58; unc-119 (+)] IV. | This study |
| JDU613 | ijmSi31 [pJD446_pJD362_Mos2_Pmex-5_mCherry_his11_3'UTRtbb-2]II; unc-119(ed3) III?; npp-13(Syb1526[gfp::npp-13])I | This study |
| JDU616 | ijmSi31 [pJD446_pJD362_Mos2_Pmex-5_mCherry_his11_3'UTRtbb-2]II; RAMP4::YFP+unc-119(e2498) III | This study |
| JDU647 | ijmSi63 [pJD520; mosII_5'mex-5_GFP::tba-2; mCherry::his-11; cb-unc-119(+)] II; baf-1(syb2768[OLLAS::mAID::baf-1])III; ieSi38 [sun-1p::TIR1::mRuby::sun-1 3'UTR + Cbr-unc-119(+)] IV | This study |
| JDU664 | ijmSi31 [pJD446_pJD362_Mos2_Pmex-5_mCherry_his11_3'UTRtbb-2]II; mel-28(bq5[GFP::mel-28]) unc-119(ed3) III? | This study |
| JDU714 | npp-7(phx1469[sgfp::npp-7])I; ijmSi31 [pJD446_pJD362_Mos2_Pmex-5_mCherry_his11_3'UTRtbb-2]II; unc-119(ed3) III? | This study |
| JDU734 | npp-12(phx3417(gfp::npp-12))I ; ijmSi31 [pJD446_pJD362_Mos2_Pmex-5_mCherry_his11_3'UTRtbb-2]II; unc-119(ed3) III? | This study |
| JDU761 | itIs37 [pie-1p::mCherry::H2B::pie-1 3'UTR + unc-119(+)] IV. orIs13 [pie-1p::GFP::npp-22::pie-1 3'UTR + unc-119(+)] ; ijmSi31 [pJD446_pJD362_Mos2_Pmex-5_mCherry_his11_3'UTRtbb-2]II; | This study |
| JDU803 | egxSi117 [pmex-5p::npp-20::gfp;;pie-1 3′UTR + unc119(+)] I; ijmSi31 [pJD446_pJD362_Mos2_Pmex-5_mCherry_his11_3'UTRtbb-2]II; unc-119(ed3) II | This study |
| JDU815 | unc-119(ed3)?, ruIs32[pAZ132; pie-1/GFP::H2B histone] III, npp-8(phx1475[mcherry::npp-8]) IV | This study |
| OD83 | qaIs3507 [pie-1::GFP::LEM-2 + unc-119(+)]; ltIs37 [(pAA64) pie-1p::mCherry::his-58 + unc-119(+)] IV. | Ródenas et al. 2009 |

| | | |
|---|---|---|
| OD270 | unc-119(ed3) III; ojls23[SP12(C34B2.10)::GFP unc-119(+)]; ltls37[pAA64; pie-1/mCHERRY::his-58; unc-119 (+)] | Penfield et al. 2020 |
| PHX2768 | baf-1(syb2768[OLLAS::mAID::baf-1])III; ieSi38 [sun-1p::TIR1::mRuby::sun-1 3'UTR + Cbr-unc-119(+)] IV | This study |
| WLP584 | jjls1092[pNUT1)npp-1::GFP+unc-119(+)]; ltls37[Ppie-1::mCherry::his-58;unc-119(+)]IV | Andy Golden et al. 2009 |
| OD999 | unc-119(ed3)III; ltSi245[pNH42; Pnpp-18::GFP-npp-18; cb-unc119(+)]II; ltls37[pAA64; pie-1/mCherry::his-58; unc-119 (+)] IV | Hattersley et al. 2016 |
| OD1496 | unc-119(ed3)III; ltSi464[pNH103; Pmex-5::npp6::GFP::tbb-2 3'UTR; cbunc-119(+)]I; ltls37[pAA64; pie-1/mCherry::his-58; unc-119 (+)] IV | Hattersley et al. 2016 |
| OD1498 | unc-119(ed3)III; ltSi465[pNH104; Pmex-5::npp-15::GFP::tbb-2 3'UTR; cb-unc-119(+)]I; ltls37[pAA64; pie-1/mCherry::his-58; unc-119 (+)] IV | Hattersley et al. 2016 |

**Supplementary table 1:** *C. elegans* strains used in this study
