## Supplementary Table 2 for "An interkinetic envelope surrounds chromosomes between meiosis I and II in C. elegans oocytes"

| Gene name | Primary sequence name | Oligonucleotide sequences | Template used | Concentration (µg/µl) |
|---|---|---|---|---|
| baf-1 | B0464.7 | AATTAACCCTCACTAAAGGCGTAGAAAATATTTTATTTGGAGG | gDNA | 2,1 |
| | | TAATACGACTCACTATAGGATGTCGACTTCTGTTAAGCATCG | | |
| knl-1 | C02F5.1 | TAATACGACTCACTATAGGttcacaaacttggaagccgctg | gDNA | 3,1 |
| | | AATTAACCCTCACTAAAGGaatctcgaatcaccgaaatgtc | | |
| lin-5 | T09A5.10 | AATTAACCCTCACTAAAGGatcgccgaggaggcacaatt | cDNA | 2,9 |
| | | TAATACGACTCACTATAGGgcttcattaccacacttgcg | | |
| mel-28 | C38D4.3 | AATTAACCCTCACTAAAGGCGTCAGCGGTTCAGTTCTTC | gDNA | 1,48 |
| | | TAATACGACTCACTATAGGCGTGTCCTGGATGTTCTGG | | |
| npp-6 | F56A3.3 | AATTAACCCTCACTAAAGGTTGGAACATGGTTGACATCG | gDNA | 1,75 |
| | | TAATACGACTCACTATAGGTTCCAAGAACTGATTGCAGG | | |
| npp-7 | T19B4.2 | AATTAACCCTCACTAAAGGAAGTCAACCAAAACCTTCCG | gDNA | 1,35 |
| | | TAATACGACTCACTATAGGAAGTTCCGTTTTGGAATTGC | | |
| npp-12 | T23H2.1 | AATTAACCCTCACTAAAGGTTGTGACTCCACTGAAACCC | gDNA | 1,42 |
| | | TAATACGACTCACTATAGGAACATTCAATGGGCACTTGG | | |
| npp-19 | R06F6.5 | AATTAACCCTCACTAAAGGTCATACTTGAGCGAGTTATTTGC | gDNA | 1,15 |
| | | TAATACGACTCACTATAGGACACCGCTAGCCCGCTCAACAC | | |
| npp-25 | Y37D8A.17 | AATTAACCCTCACTAAAGGATGATCATCAATCTTCATCC | cDNA | 2,35 |
| | | TAATACGACTCACTATAGGATGACTGGCGGAGCTCTAGC | | |
| npp-15 | C29E4.4 | AATTAACCCTCACTAAAGGTTCCTAAGTTCCCTATTTGG | cDNA | 1,97 |
| | | TAATACGACTCACTATAGGAATAGTTGCAGAATTTTCCG | | |
| VRK-1 | F28B12.3 | TAATACGACTCACTATAGGaacgaatcatctgccgtcgttgg | gDNA | 2,4 |
| | | AATTAACCCTCACTAAAGGctctttgataatcatcgatcg | | |

**Supplementary table 2:** dsRNA used in this study
