## Supplementary Table 3 for "An interkinetic envelope surrounds chromosomes between meiosis I and II in C. elegans oocytes"

| Figure | Antibody | Source | Concentration |
|---|---|---|---|
| 1F | rabbit anti-LEM-2 | Novus Biologicals 48540002 | 1/1000 |
| | mouse anti-α-tubulin (FITC-labeled DM1α) | Sigma-Aldrich F2168 | 1/200 |
| | Dylight 549-conjugated goat anti-rabbit | Jackson ImmunoResearch 111-165-144 | 1/100 |
| S1B | mouse anti-GFP | Roche 11814460001 | 1/200 |
| | human anti-α-tubulin | Institut Curie, Paris, France | 1/200 |
| | rabbit anti-CHMP-7 | Gift from Anjon Audhya | 1/1000 |
| | rabbit anti-VPS-32 | Gift from Anjon Audhya | 1/1000 |
| | Dylight 461-conjugated goat anti-mouse | Jackson ImmunoResearch 115-225-071 | 1/100 |
| | Dylight 549-conjugated donkey anti-human | Jackson ImmunoResearch 709-165-149 | 1/100 |
| | Dylight 649-conjugated goat anti-rabbit | Jackson ImmunoResearch 111-175-144 | 1/100 |
| S2A | rat anti-ollas | Novus Biologicals NBP1-06713 | 1/500 |
| | rabbit anti-LEM-2 | Novus Biologicals 48540002 | 1/1000 |
| | mouse anti-a-tubulin (FITC-labeled DM1α) | Sigma-Aldrich F2168 | 1/200 |
| | Dylight 649-conjugated Donkey anti-rat | Jackson ImmunoResearch 112-175-143 | 1/100 |
| | Dylight 549-conjugated goat anti-rabbit | Jackson ImmunoResearch 111-165-144 | 1/100 |

**Supplementary table 3:** antibodies used in this study
